# Stable isotopes disentangle niche partitioning and species co-occurrence in a multi-level marine mutualism

**DOI:** 10.1101/2023.11.22.568307

**Authors:** Benjamin M. Titus, Catheline Y. M. Froehlich, Clayton Vondriska, Ronald Baker, Eleanor M. Caves

## Abstract

Ecologists have long sought general explanations for the co-occurrence of ecologically similar taxa. Niche theory explains co-occurrence via functional differences among taxa that reduce competition and promote resource partitioning. Alternatively, the unified neutral theory of biodiversity and biogeography suggests that co-occurrence can be attributed to stochastic processes, and thus, presupposes that ecologically similar species that occur in sympatry are functionally analogous. We test these alternative hypotheses in multiple dimensions using the most diverse crustacean-sea anemone symbiosis from coral reefs in the Tropical Western Atlantic. δ^13^C and δ^15^N stable isotope analyses of six crustacean symbionts that co-occur around the host anemone *Bartholomea annulata* exhibit highly differentiated isotopic niche space spanning three trophic levels. As multiple crustacean species within the symbiosis have been documented as cleaners that remove parasites from reef fishes, we extended our investigation into the broader cleaner community. Our stable isotope analyses of cleaners shows that only Pederson’s cleaner shrimp *Ancylomenes pedersoni* exhibits δ^15^N isotopic signatures that are consistent with a dedicated cleaning lifestyle. Co-occurring species that have been previously described to clean reef fishes such as *Periclimenes yucatanicus, Stenopus hispidus,* and *Stenorhynchus seticornis* all occupy trophic levels well below *An. pedersoni*. Taken together, our data are consistent with the expectations of niche theory: co-occurring symbiotic crustaceans have highly partitioned niche space with low levels of functional redundancy. Finally, our findings reinforce and extend the ecological importance of *An. pedersoni* as likely the only dedicated cleaner shrimp on coral reefs in the Tropical Western Atlantic.

## Introduction

Within an ecological community species coexistence has classically been explained by niche theory (e.g. Hutchison 1957, Vandermeer 1972, Colwell and Rangel 2009, Koffel et al. 2021), which posits that functional differences among taxa reduce interspecific competition and promote resource partitioning and co-occurrence. Under this framework ecological communities are not stochastic assemblages of functionally redundant taxa but represent the combined outcome of deterministic, selective processes. Alternatively, the unified neutral theory of biodiversity and biogeography (Hubbell 2001, Rosindell et al. 2012) suggests that ecological communities and species co-occurrence can be attributed to stochastic processes, and that diversity within an ecological community does not need to be explained by invoking species’ functional differences. Under this framework more species co-exist within an assemblage than the number of niches available, and co-existence of ecologically similar species can be attributed to a balance of demographic processes such as speciation, extinction, and immigration at various biogeographic scales.

The diverse mutualisms found on tropical coral reefs provide ideal frameworks to test these competing hypotheses (Dornelas et al. 2006, Rosindell et al. 2012, Connolly et al. 2017). Coral reefs contain 25% of all marine biodiversity on <0.1% of the sea floor and competition for space and resources is famously intense (e.g. Spalding and Grenfell 1997, McCook et al. 2001, Chadwick and Morrow 2012, Bonin et al. 2015). Many reef-dwelling taxa have thus evolved tightly linked symbioses as novel ways to acquire food and space (Castro 1988). These include nutritional endosymbioses with dinoflagellates, host-symbiont shelter symbioses (e.g. clownfish and sea anemones), and cooperative cleaning symbioses, among others (e.g. Muscatine and Porter 1977, Castro 1988, Pochon and Pawlowski 2006, Vaughan et al. 2017, Feeney et al. 2019, Caves 2021). Typically, a single reef harbors many mutualistic species that are ecologically similar and appear functionally redundant. For example, multiple species of cleaner fishes and cleaner shrimps co-occur on a given reef and can even provide cleaning services to the same reef fish species (e.g. Titus et al. 2015a, Rose et al 2020). However, the processes that allow for this degree of co-occurring functional diversity remain largely unexplored for most systems.

On coral reefs in the Tropical Western Atlantic (TWA), large sea anemones (Cnidaria: Anthozoa: Actiniaria) form the hub of diverse multi-level mutualisms (Briones-Fourzán et al. 2012, Cantrell et al. 2015, Huebner et al. 2012a, 2019, Titus et al. 2015a, 2017a, 2019, Caves et al. 2018). Sea anemones in the TWA serve as hosts to a diverse assemblage of over 15 crustacean species, many of which belong to lineages that have independently evolved symbiotic lifestyles (Briones-Fourzán et al. 2012, Colombara et al. 2017, Titus and Daly 2017, Horka et al. 2018, Brooker et al. 2019, Huebner et al. 2019). While some of these crustaceans are highly specialized obligate mutualists on a single host anemone species, others are obligate symbionts of sea anemones broadly but retain host species flexibility, and some are non-obligate, facultative anemone dwellers (Briones-Fourzán et al. 2012, Titus and Daly 2017, Huebner et al. 2019). Rates of co-occurrence are high among crustacean ectosymbionts- up to seven species have been observed occupying a single host simultaneously (Briones-Fourzán et al. 2012, Colombara et al. 2017, Brooker et al. 2019, Huebner et al. 2019), and intraspecific groups sizes can vary from solitary individuals to monogamous pairs to cohorts of up to 10 or more conspecifics. Thus, dozens of individuals across multiple species can co-occur on one anemone host, and because of this, crustacean symbionts often partition the microhabitat around anemones (Briones-Fourzán et al. 2012, Colombara et al. 2017, Brooker et al. 2019, Huebner et al. 2019). It has been hypothesized that microhabitat partitioning reduces competition among co-occurring anemone symbionts (Huebner et al. 2019). However, more symbiotic species can exist than discrete microhabitat zones around a host anemone, and the ecological similarity among members of the symbiont community can be high. It remains unclear if the fine-scale spatial partitioning of mere centimeters surrounding a host is reflective of broader underlying niche partitioning that results from unknown functional differences (e.g. diet) among the crustacean symbionts.

Of particular interest are the functional roles of multiple crustacean symbionts that have been observed cleaning reef fishes (McCammon et al. 2010, Medeiros et al. 2011, Huebner et al. 2012a, b, Titus et al. 2015a, 2015b, 2017a, 2017c, 2019, Caves et al. 2018, Rose et al. 2020). Cleaning symbioses are ubiquitous and ecologically important mutualisms on coral reefs where client reef fish pose at a cleaning station (often a discrete habitat marked by a sea anemone, coral, or prominent location on the reef) and have ectoparasites removed by a smaller cleaner species- typically a cleaner shrimp or cleaner fish (Côté 2000, Vaughan et al. 2017, Caves 2021). Cleaning symbioses have evolved many times independently within decapod crustaceans (Vaughan et al. 2017, Horka et al. 2018). Some species are described as “dedicated” cleaners, meaning they are committed to a cleaning lifestyle for all their non-larval ontogeny, while others are facultative cleaners and only clean occasionally or during specific life stages (Vaughan et al. 2017). For many species, cleaner status is based solely off anecdotal evidence and one-off observations without in-depth investigation into their behavior, diet, or ecology, thus, it has been suggested that the “cleaner” label is overused when describing the functional role of many putative cleaner species (Vaughan et al. 2017). Recently, De Grave et al (2021) used stable isotopes to demonstrate that, amongst a group of symbiotic palaemonid shrimps and their invertebrate hosts in Japan, the shrimp species with the highest nitrogen ratio was a known cleaner. Dedicated cleaners should have high trophic levels since they consume parasites from reef fishes, and thus, stable isotopes may be a valuable tool in helping to distinguish true cleaners from facultative or non-cleaners.

Here we use a multi-level sea anemone symbiosis from the TWA and stable isotope analyses to test alternative hypotheses surrounding species co-occurrence and niche partitioning. Stable isotopes represent a powerful approach to disentangle dietary niche partitioning, and give insight into the functional roles of organisms (which is especially valuable in crustaceans, where behavioral observations are limited) since measures of isotopic niche space reflect a long-term dietary mean. We focus on the corkscrew sea anemone *Bartholomea annulata*, the most abundant host anemone in the TWA (Briones-Fourzán et al. 2012, Titus et al. 2017b, O’Reilly et al. 2018), and a suite of crustacean symbionts that commonly co-occur on *B. annulata* (Table 1, Fig. 1). Prior research and observations on this symbiosis provide support for both niche theory and neutral theory, meaning open questions remain regarding how organisms in this system manage to coexist. In support of predictions made by niche theory, host sea anemones are rare and do not obtain high densities on coral reefs (Briones-Fourzán et al. 2012, Titus et al. 2017b, O’Reilly et al. 2018, Huebner et al. 2019). Resource limitation is expected to increase competitive interactions among co-occurring species and lead to greater niche partitioning. Microhabitat partitioning around *B. annulata* has been well documented within this symbiosis (Huebner et al. 2019), which has been hypothesized as a potential mechanism to reduce competition. Alternatively, in support of predictions made by the unified neutral theory, not all anemones are occupied by crustacean symbionts (Briones-Fourzán et al. 2012, Huebner et al. 2019), and so the degree to which crustacean symbionts may compete for space isn’t immediately obvious. Species specific demographic processes, rather than interspecific competition for food, may instead be responsible for limiting population sizes and driving the variation in abundance that is observed for each species. Finally, many species are ecologically similar, suggesting some degree of overlapping functional redundancy within the symbiotic crustacean community.

**Fig. 1.**
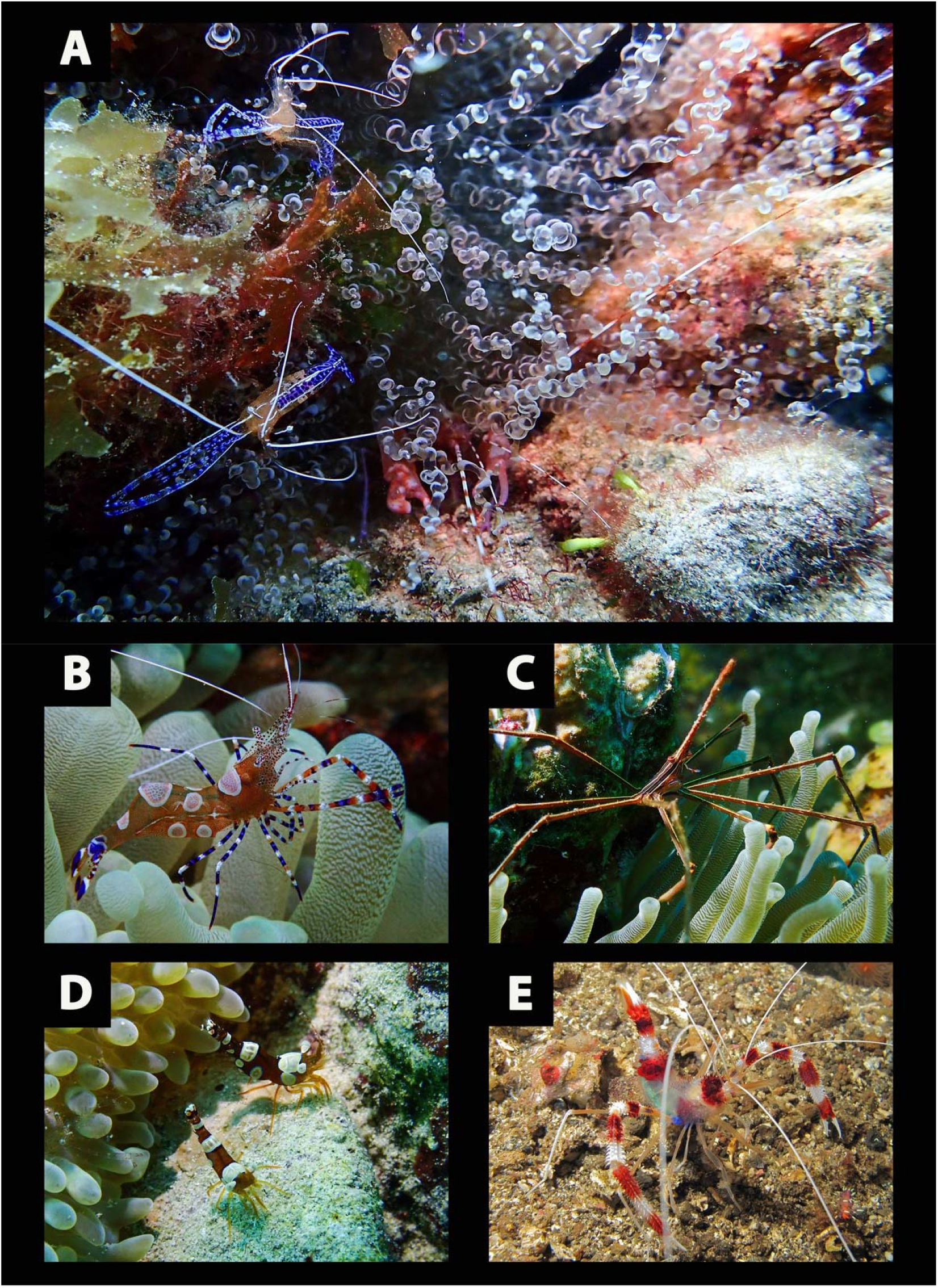
Focal taxa: A) Corkscrew sea anemone *Bartholomea annulata* (center, spiral shaped tentacles visible) with Pederson’s cleaner shrimp *Ancylomenes pedersoni* (purple shrimp, white antennae) and pistol snapping shrimp in the *Alpheus armatus* species complex (red shrimp, long red and white antennae visible). Snapping shrimps in genus *Alpheus* live under host *B. annulata* while *A. pedersoni* are mostly found on peripheral locations. B) Spotted cleaner shrimp *Periclimenes yucatanicus* on the tentacles of the Giant Caribbean sea anemone *Condylactis gigantea*. C) Yellowline arrow crab *Stenorhynchus seticornis*. D) Caribbean sexy shrimp *Thor dicaprio*. E) Banded Coral Shrimp and putative cleaner species *Stenopus hispidus.* Photo E by Bernard Dupont used under creative commons license CC BY-SA 2.0

**Table 1.**
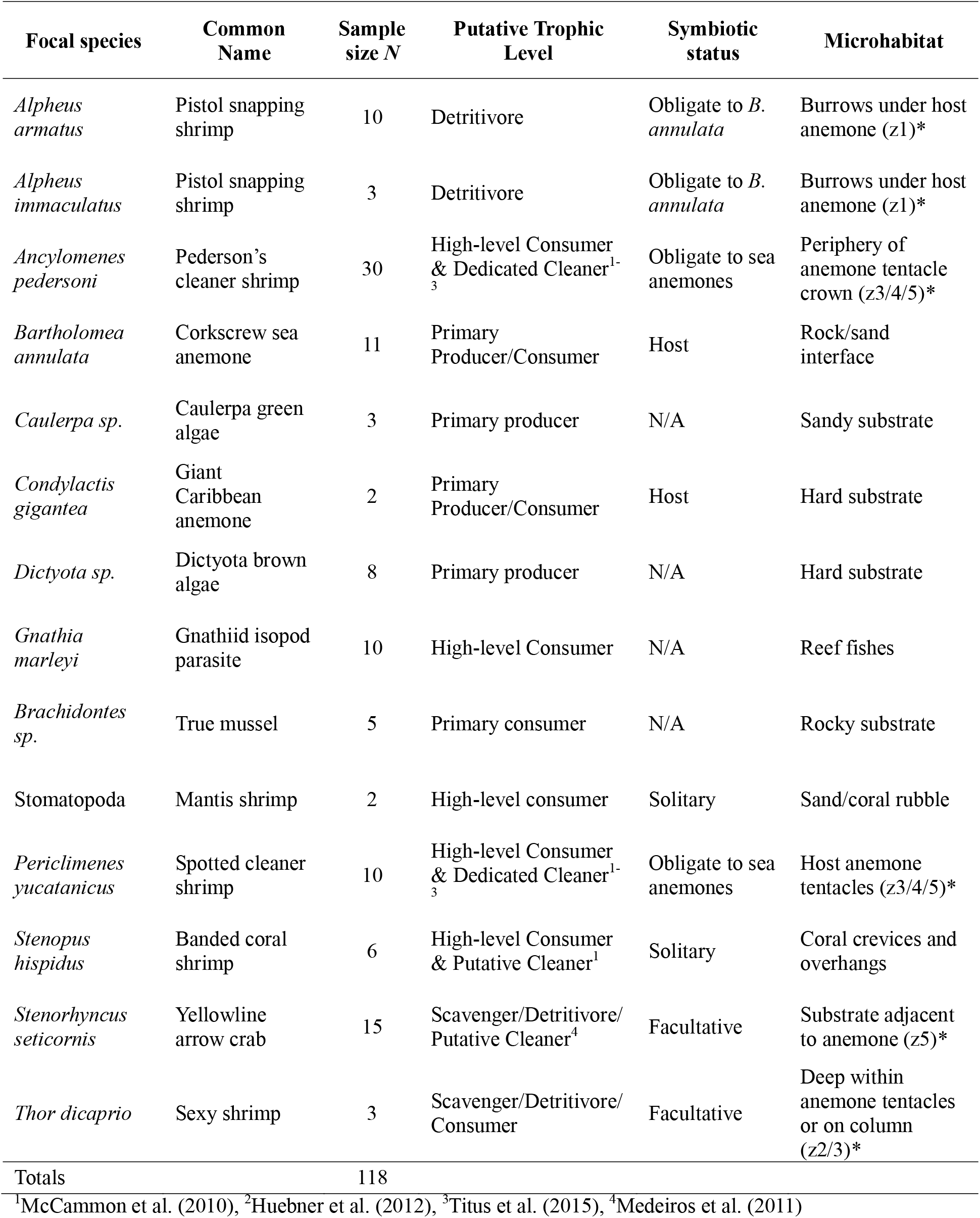
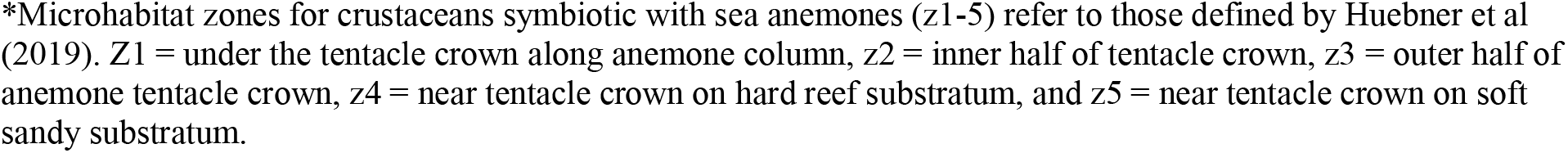
Focal taxa included in this study collected from St. Thomas, US Virgin Islands along with sample sizes (N), putative trophic levels, symbiotic status with host sea anemones, and microhabitats.

| Focal species | Common Name | Sample size *N* | Putative Trophic Level | Symbiotic status | Microhabitat |
| --- | --- | --- | --- | --- | --- |
| *Alpheus armatus* | Pistol snapping shrimp | 10 | Detritivore | Obligate to *B. annulata* | Burrows under host anemone (z1)* |
| *Alpheus immaculatus* | Pistol snapping shrimp | 3 | Detritivore | Obligate to *B. annulata* | Burrows under host anemone (z1)* |
| *Ancylomenes pedersoni* | Pederson's cleaner shrimp | 30 | High-level Consumer & Dedicated Cleaner<sup>1-3</sup> | Obligate to sea anemones | Periphery of anemone tentacle crown (z3/4/5)* |
| *Bartholomea annulata* | Corkscrew sea anemone | 11 | Primary Producer/Consumer | Host | Rock/sand interface |
| *Caulerpa sp.* | Caulerpa green algae | 3 | Primary producer | N/A | Sandy substrate |
| *Condylactis gigantea* | Giant Caribbean anemone | 2 | Primary Producer/Consumer | Host | Hard substrate |
| *Dictyota sp.* | Dictyota brown algae | 8 | Primary producer | N/A | Hard substrate |
| *Gnathia marleyi* | Gnathiid isopod parasite | 10 | High-level Consumer | N/A | Reef fishes |
| *Brachidontes sp.* | True mussel | 5 | Primary consumer | N/A | Rocky substrate |
| Stomatopoda | Mantis shrimp | 2 | High-level consumer | Solitary | Sand/coral rubble |
| *Periclimenes yucatanicus* | Spotted cleaner shrimp | 10 | High-level Consumer & Dedicated Cleaner<sup>1-3</sup> | Obligate to sea anemones | Host anemone tentacles (z3/4/5)* |
| *Stenopus hispidus* | Banded coral shrimp | 6 | High-level Consumer & Putative Cleaner<sup>1</sup> | Solitary | Coral crevices and overhangs |
| *Stenorhynchus seticornis* | Yellowline arrow crab | 15 | Scavenger/Detritivore/ Putative Cleaner<sup>4</sup> | Facultative | Substrate adjacent to anemone (z5)* |
| *Thor dicaprio* | Sexy shrimp | 3 | Scavenger/Detritivore/ Consumer | Facultative | Deep within anemone tentacles or on column (z2/3)* |
| Totals |  | 118 |  |  |  |

<sup>1</sup>McCammon et al. (2010), <sup>2</sup>Huebner et al. (2012), <sup>3</sup>Titus et al. (2015), <sup>4</sup>Medeiros et al. (2011)
\*Microhabitat zones for crustaceans symbiotic with sea anemones (z1-5) refer to those defined by Huebner et al (2019). Z1 = under the tentacle crown along anemone column, z2 = inner half of tentacle crown, z3 = outer half of anemone tentacle crown, z4 = near tentacle crown on hard reef substratum, and z5 = near tentacle crown on soft sandy substratum.

We test the null hypothesis established by the unified theory of biodiversity and biogeography that ecologically similar co-occurring species are functionally redundant, and by extension, occupy overlapping isotopic niche and trophic space. We use two communities with partially shared membership: 1) the sea anemone-crustacean symbiosis, which includes the host anemones *B. annulata* and *Condylactis gigantea*, along with crustacean symbionts *Alpheus armatus, Al. immaculatus*, *Ancylomenes pedersoni, Periclimenes yucatanicus, Stenorhynchus seticornis,* and *Thor dicaprio*, and 2) the community of crustaceans that serve as cleaners, which includes the dedicated cleaner species *An. pedersoni* and *P. yucatanicus,* as well as putative cleaner species *Stenopus hispidus* and *S. seticornis.* For both datasets, non-overlapping isotopic niche space partitioning would thus provide compelling evidence that co-occurring species are not functionally redundant and are acquiring resources in different ways.

## Methods

### Study sites, focal taxa, and sample collection

Our study was conducted on three fringing reefs surrounding St. Thomas, US Virgin Islands: 1) Brewers Bay (18°20’27.95”N, 64°58’42.41”W), a shallow (∼3-6m depth) inshore fringing reef adjacent to the Maclean Marine Science Center at the University of the Virgin Islands, 2) Black Point (18° 20’ 40.30”N, 64° 59’ 6.67” W), a shallow (∼3-7m depth) inshore fringing reef at the western tip of Brewers Bay less than 1km from our Brewers Bay sampling locality, and 3) Flat Cay (18°19’1.77”N, 64°59’7.49”W), an offshore fringing reef ∼2km from Brewers Bay that is characterized by slightly greater depth (∼7-12m depth) and more oligotrophic conditions (i.e. clearer water, lower nutrients). We chose these study localities because St. Thomas, USVI, and these reef sites specifically, are among the most heavily studied in the TWA for crustacean-sea anemone symbioses (McCammon et al. 2010, Huebner et al. 2012a, b, 2019, O’Reilly et al. 2018). These same localities have also been used previously to show how crustacean symbionts spatially partition microhabitat space around sea anemones (Huebner et al. 2019).

At each reef, focal taxa (Table 1, Fig. 1) were collected by hand using SCUBA and were transported to the University of the Virgin Islands for processing. The host sea anemone *Bartholomea annulata* (Fig. 1a) was sampled by collecting 3-5 tentacles per individual using forceps. Host anemones, which represent a potential food source for their symbiotic crustaceans (e.g. Suzuki and Kayashi 1997, Fautin et al. 1995, Khan et al. 2004), partition the bulk of their endosymbiotic algae in their tentacles, and the majority of their energetic requirements are met via photosynthetically derived sources. Thus, any tissue ingested by crustacean symbionts would represent a combination of host and dinoflagellate material, and by extension, acquire an isotopic signature more similar to a primary rather than secondary consumer. Other host anemone species are rare at our field sites (Huebner et al. 2019), but *Condylactis gigantea* were sampled opportunistically and included in our stable isotope analyses as comparative samples (Table 1, Fig. 1b).

At each host anemone, we also sampled the available symbiotic crustacean community by hand using whirl-pak bags. Our efforts focused on, and reflect, the most common crustacean symbionts found in association with *B. annulata*. These include two sister species of snapping shrimps that are obligate to *B. annulata*: *Alpheus armatus* and *A. immaculatus* (Table 1, Fig. 1a). Both species are part of the broader *Alpheus armatus* species complex (Hurt et al. 2013) and were collected using clove oil to temporarily stun the animals, which were then placed into a whirl-pak bag and revived with ambient seawater. Both snapping shrimp species burrow under the host anemone and have been shown to be important in protecting the anemone from predatory fireworms as well as removing sediment that could smother their host (McCammon and Brooks 2014, Perez-Botello et al. 2021). Although these species have been observed to feed on detritus (Knowlton 1980), their trophic level is unverified.

Next, we collected two closely related species of palaemonid shrimps that are obligate symbionts of sea anemones but that have retained flexibility in their host associations (Table 1, Fig. 1): Pederson’s cleaner shrimp, *Ancylomenes pedersoni* (Fig. 1a), and the spotted cleaner shrimp, *Periclimenes yucatanicus* Fig. 1b). At our field sites in St. Thomas, where *C. gigantea* is rare (Huebner et al. 2019), *P. yucatanicus* is primarily associated with *B. annulata* and often co-occurs with *An. pedersoni*. When both species co-occur on the same individual host, *P. yucatanicus* occupies central microhabitat locations on the sea anemone tentacles while *An. pedersoni* occupies more peripheral locations on or around the tentacles (Huebner et al. 2019).

Both *An. pedersoni* and *P. yucatanicus* have been described as cleaner shrimps that engage in a mutualism by which they remove parasites from reef fishes (Huebner et al. 2012a, b, Titus et al. 2015a, 2015b, 2017a, 2017c 2019, Caves et al. 2018, Rose et al. 2020). The ecology and behavior of *An. pedersoni* has been studied extensively both in St. Thomas and elsewhere, and it is considered a dedicated cleaner shrimp (Huebner et al. 2012a, 2012b, Titus et al. 2017a, 2019, Caves et al. 2018, Romain et al. 2020, McCloskey et al. 2023), known to clean over 50 species of reef fish clients from over 23 families, and shown to be effective at reducing parasite loads for one common reef fish parasite (McCammon et al 2010, Titus et al. 2015a, 2015b 2019, Romain et al. 2020). We expect dedicated cleaners to occupy higher-level consumer trophic levels in stable isotope analyses, although it remains unknown whether *An. pedersoni* supplements its diet with other food resources. The cleaner status of *P. yucatanicus*, however, is more equivocal. Titus et al. (2017a) demonstrated that while *P. yucatanicus* engages in true cleaning interactions with reef fish, it cleans less frequently, engages in shorter cleaning bouts, and has fewer client visits and less client species than *A. pedersoni*. McCammon et al. (2010) showed *P. yucatanicus* did not significantly reduce parasite loads for a common reef parasite in St. Thomas, USVI, but did reduce mean parasite size. This species is consistently listed as a dedicated cleaner species in the broader literature (e.g. Vaughan et al. 2017, Horka et al. 2018), but its ecological function as a cleaner remains ambiguous.

We further collected two common anemone associates with less strict host specificities to *B. annulata*, the Caribbean “sexy shrimp” or “squat anemone shrimp” *Thor dicaprio* (recently differentiated from the previously recognized circumtropical *T. amboinensis* via molecular and morphological evidence, Titus et al. 2018, Anker and Baeza 2021), and the yellowline arrow crab *Stenorhynchus seticornis* (Table 1, Fig. 1)*. T. dicaprio* can be found on a broad range of hosts in the TWA including anemones, corals, corallimorpharians, and crinoids (Bruce 1976, Criales 1984, Khan et al. 2004, Hoeksema and Fransen 2011, Briones-Fourzan et al. 2012). When in association with *B. annulata*, *T. dicaprio* mostly resides under anemone tentacles or on the column of the host (Huebner et al. 2019). The ecological role and trophic level of *T. dicaprio* are unknown, but it is commonly collected in the ornamental aquarium trade and fed an omnivorous diet in captivity (Bartilotti et al. 2016). The yellowline arrow crab *S. seticornis* is a facultative symbiont of anemones, as it also associates with a number of benthic invertebrates and can be found free living in cracks and crevices on coral reefs (Hays et al. 1998, Briones-Fourzan et al. 2012, Huebner et al. 2019). When in association with sea anemones this species is found on the margins of anemone tentacles (Huebner et al. 2019). The trophic level and functional role of this species is unknown, but many crab species feed on detritus. Interestingly, in Brazil Medeiros et al. (2011) observed this species engaging in cleaning interactions with at least five species of reef fish and we have observed cleaning by arrow crabs on at least two occasions in Curacao, Netherlands Antilles (E. Caves, pers. obs). Thus, at least some evidence suggests it may be a facultative cleaner.

Lastly, we collected the banded coral shrimp *Stenopus hispidus* (Table 1, Fig. 1). *S. hispidus* is a putative cleaner shrimp (Limbaugh 1961, Sazima et al. 2004, Vaughan et al. 2017), but is free-living and does not associate with sea anemones. Like *P. yucatanicus* and *S. seticornis* its true ecological function as a cleaner species is uncertain and documentation as a cleaner limited to anecdotal observations (Limbaugh 1961). McCammon et al. (2010) showed that it had no significant effect on monogenean flatworm abundance on Caribbean Blue Tang, but did show a significant effect on average parasite size. Although not an associate of *B. annulata*, we collected *S. hispidus* from crevices and reef overhangs at Brewers Bay, given their putative cleaner status.

To compare the trophic niche spaces our focal taxa above we collected control samples of primary producers, primary consumers, and secondary consumers to represent key functional groups within Brewers Bay as this was where most of our samples were collected. To capture benthic primary production we collected green algae samples from the genus *Caulerpa* and brown algae samples from the genus *Dictyota*. To capture planktonic production and the primary consumer community we collected mussels from the genus *Brachidontes*. Similarly, to capture the higher-level consumer community we collected mantis shrimps (Stomatopoda) from the “smasher” guild that use their blunt dactyl claws to smash open hard-shelled prey, primarily crustaceans and gastropods. Finally, to help characterize the cleaner community and differentiate dedicated from occasional cleaner species that might supplement their diets elsewhere, we included final stage juvenile gnathiid isopod parasites (*Gnathia marleyi*) in our analyses. Gnathiid isopods are common reef fish ectoparasites that feed on the blood and lymph of over 80 species of reef fishes in the Caribbean during their juvenile life stages. The gnathiid samples used here had been collected previously and were being maintained in a mesocosm facility by feeding on a mix of damselfish species in the genus *Stegastes.* These gnathiids had parasitized their final host fish in captivity 2-3 days before collection and inclusion in this study.

### Stable isotope analysis

Following field collection, all samples were transported to the Environmental Analysis Laboratory at the University of the Virgin Islands where they were immediately euthanized in a seawater ice bath. Abdominal muscle tissue from large (>2cm) crustacean samples was dissected, then all samples were soaked in distilled water for >30 min to remove salts and dried for 24hrs at 70°C. Samples were then transported to the University of California at Santa Barbara (UCSB) Marine Science Institute Analytical Lab where dried tissue was ground into a fine powder with mortar and pestle and 500-700ug of tissue was packed into tin capsules. We did not acidify our samples prior to stable isotope analysis. Although this practice is common, acidification aimed at removing inorganic carbonates can have large effects on nitrogen ratios (Vafeidou et al. 2013). Additionally, dissected muscle tissue should be free of carbonates, crustacean exoskeletons should have minimal carbonate composition, and acidifying whole samples has been shown to have no significant effect on carbon ratios within Malacostraca (Mateo et al. 2008). Stable isotope ratios of carbon (^13^C/^12^C) and nitrogen (^15^N/^14^N) were measured using a Thermo Finnigan Delta-Plus Advantage isotope mass spectrometer coupled with a Costech EAS elemental analyzer. Instrument calibration was conducted using acetanilide reference standards run at the beginning of each set of 35 samples and tested every 5 samples within each set. Instrument precision, determined using replicate analyses of L-glutamic acid USGS40, was ± 0.02 and ± 0.09 for ^15^N and ^13^C, respectively. Stable isotope ratio values are denoted in parts per thousand (‰) using the standard δ notation, where δX = [R_SAMPLE_/R_STANDARD_ – 1) x 1000], with R being the ratio of heavy to light isotopes. The standard reference material was Pee Dee Belemite (vPDB) and air for carbon and nitrogen analyses respectively.

Prior to downstream analyses, Carbon isotopic ratios (δ^13^C) were lipid corrected for invertebrate species whose carbon to nitrogen ratios were above 4, using the lipid-normalizing model introduced by Kijunen et al. (2006). We then used these values to calculate the isotopic niche space and trophic level for each species. Isotopic niche space is defined here as a 2- dimensional multivariate space where the axes represent the composition of δ^13^C and δ^15^N in an animal’s tissue following Newsome et al., (2007). Trophic level was calculated using δ^15^N only. We tested for overlap in both dimensions.

We visualized isotopic niche space for the full dataset with biplots for all samples individually and by calculating the mean ± SE for each species. The unified theory of biodiversity and biogeography predicts that ecologically similar co-occurring species are functionally redundant, and by extension, occupy overlapping isotopic niche and trophic space. Using both isotopic ratios (δ^13^C and δ^15^N) we tested the null hypothesis established by the unified theory using two communities with partially shared membership: 1) the anemone-crustacean symbiosis, which includes the host anemones *B. annulata* and *C. gigantea*, along with crustacean symbionts *Al. armatus, Al. immaculatus*, *An. pedersoni, P. yucatanicus, S, seticornis,* and *T, dicaprio*, and 2) the broader crustacean cleaner community, which includes the dedicated cleaner species *An. Pedersoni* and *P. yucatanicus,* as well as putative cleaner species *S. hispidus* and *S, seticornis.* We further include the gnathiid isopod *G. marleyi* as a primary reef fish parasite and potential target food source for cleaners. For both datasets, non-overlapping isotopic niche space partitioning would thus provide compelling evidence that co-occurring species are not functionally redundant and are acquiring resources in different ways. Based on De Grave et al. (2021) we expect dedicated cleaners to occupy high-level trophic levels, and specifically, secondary or tertiary consumer levels with δ^15^N ratios above reef fish parasites.

For each dataset, we calculated the isotopic niche space in R v4.2.2 (R Core Team 2022) by fitting standard ellipses (40% confidence level) to individually plotted data using the SIBER package (Jackson et al. 2011). The standard ellipse area (SEA) was fitted with Bayesian models (10^4^ iterations) and were corrected for small sample size (SEA_c_). The area of ellipse overlap between species represents the degree of overlap in isotopic niche space, which was quantified using both SEA_C_ and the percentage of ellipse area overlap. Analyses were conducted for both datasets. Differences in SEA_C_ were considered significant when ≥ 95% of posterior draws for one species was smaller than the other. Species with a percent overlap >60% were considered to exhibit significant shared isotopic niche space as per Schoener (1968). For standard ellipse and overlap calculations we pooled *C. gigantea* data with *B. annulata* data due to inadequate samples sizes needed to calculate a standard ellipse for *C. gigantea* alone, and due to the broadly similar isotopic trophic level both species appeared to occupy (Fig. 2, Fig. S1).

**Fig. 2.**
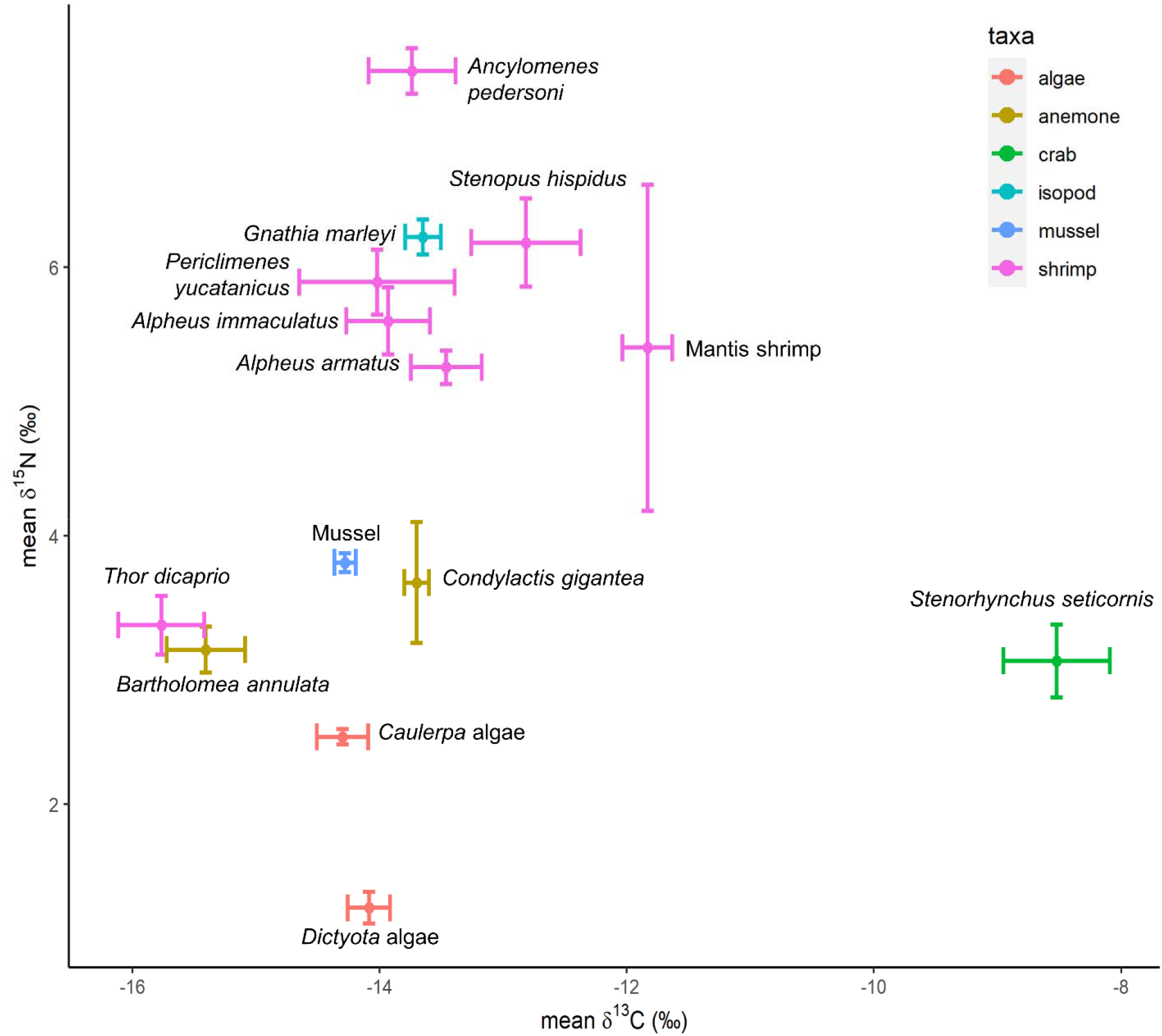
δ^13^C and δ^15^N isotopic signatures of for all species collected for this study (e.g. host sea anemones, focal crustaceans, and producers and consumers) from St. Thomas, US Virgin Islands. Data are presented as means and standard error of carbon and nitrogen values. Data are colored by broad taxonomic group and labeled by species. Vector denotes trophic level discrimination for δ^13^C (+0.39) and δ^15^N (+3.4) on x and y axes respectively.

Trophic level for each species was calculated using the package tRophicLevel (Quezada-Romegialli et al. 2018) with the following parameters: trophic discrimination factors of ΔN (3.4 ± 0.98 sd) and ΔC (0.39 ± 1.3 sd) based on Post’s (2002) assumptions 20^4^ model iterations as additional iterations did not change the variation, and oneBaseline approach using algae sources (*Dictyota* and *Caulerpa* algae) as the baseline for trophic level. Significant differences among species (fixed factor) in trophic leveling and δ^13^C isotopic niche space were tested for separately using linear models. For each linear model, data were transformed to meet normality and homoscedasticity using Q-Q plot visualization, histograms, and residuals over fitted plots, and outliers were removed if their standard residuals fell outside of 2.5 standard deviation from 0. Pairwise comparisons were made using Tukey’s *post hoc* tests to test for significant differences in trophic level and δ^13^C isotopic niche space among species. Additional R packages used to complete the analyses included: tidyverse (Wickham et al. 2019), GGally (Schloerke et al. 2022), reshape2 (Wickham 2007), plyr (Wickham 2011), lme4 (Bates et al. 2015), emmeans (Lenth et al. 2020), piecewiseSEM (Lefcheck et al. 2016), LMERConvenienceFunctions (Tremblay and Ransijn 2020).

Finally, we found that *An. Pedersoni* and *P. yucatanicus*—both obligate anemone symbionts and cleaner shrimps that were collected from both Brewers Bay and Flat Cay— appeared to exhibit a broad range of isotopic ratios. Therefore, we further explored variation in δ^13^C and δ^15^N ratios within and between species across our three sample localities at USVI. All analyses and statistical comparisons were conducted as above, and localities were included in the analyses as fixed factors.

## Results

Isotopic biplots for all data showed partitioning across both δ^13^C and δ^15^N axes (Fig. 2, Fig. S1). δ^15^N ratios for environmental control samples reflected our expected trophic levels: primary producers (algae) contained the lowest δ^15^N values, followed by the primary consumer mussels, and the higher-level consumer mantis shrimps and gnathiid isopod fish parasites had δ^15^N ratios reflective of their predatory lifestyle (Fig. 2). Filter feeding mussel samples exhibited depleted δ^13^C values, consistent with the expectation of a species acquiring carbon from pelagic primary sources.

### Crustacean symbiont niche partitioning

Among our focal taxa our data show a strong signature of isotopic niche and trophic partitioning (Fig.s 3 and 4). Within the sea anemone-crustacean symbiosis all species exhibited significantly partitioned isotopic niche space with very little overlap among co-occurring symbionts (Fig. 3a, Table S1, Fig. S2). Host anemone species had low δ^15^N ratios and low trophic levels on par with primary consumers (Fig. 3a, Fig. 4a), which is expected given their endosymbiosis with photosynthetic dinoflagellates. Crustacean symbionts with facultative symbioses with sea anemones, *T. dicaprio* and *S. seticornis*, also exhibited low δ^15^N ratios and trophic levels (Fig. 3a, Fig. 4a), comparable to the host anemones and primary consumers, but with very different δ^13^C ratios (Fig. 3a, Fig. S3). The yellowline arrow crab *S. seticornis* exhibited the most enriched δ^13^C ratios of all species (Fig. 3a, Fig. S3), a signature typically associated with benthic carbon sources, but did not share isotopic niche space with any other species within the symbiosis (Fig. 3a, Table S1, Fig. S2). Samples of *T. dicaprio* were more carbon depleted, potentially a signature derived from more pelagic-based carbon sources (zooplankton/particulate organic matter, Fig. 3a, Fig. S3). Snapping shrimps *A. armatus* and *A. immaculatus*, both obligates to *B. annulata* that occupy the same anemone microhabitat, had the greatest isotopic niche overlap of any species comparisons within the symbiosis (20%) but remained well short of the 60% overlap threshold to conclude that they occupy the same isotopic niche space (Fig. 3a, Table S1, Fig. S2).

**Fig. 3.**
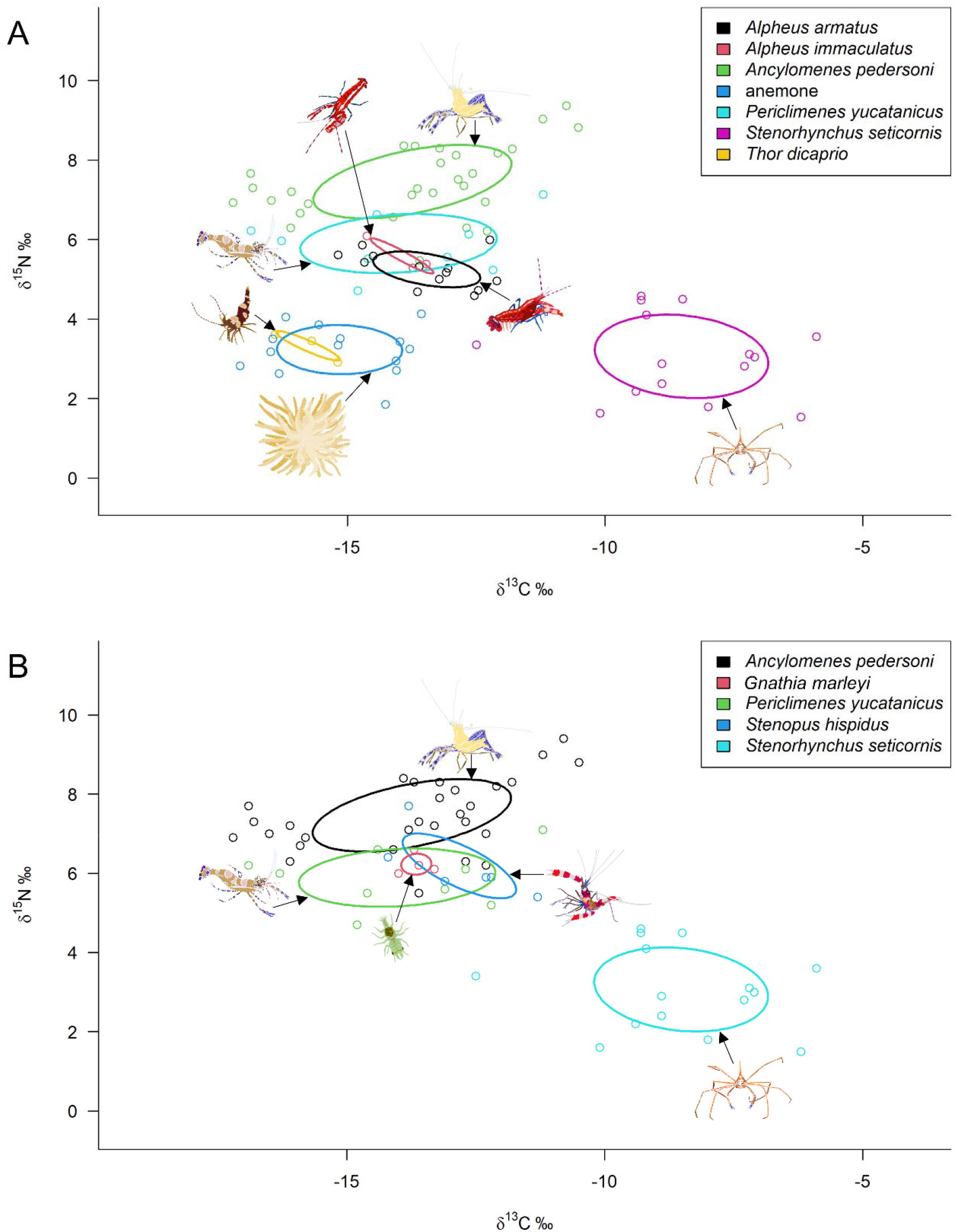
A) Isotopic Niche partitioning for the sea anemone-crustacean symbiosis and B) isotopic Niche overlap for putative crustacean cleaner species. 40% Standard Ellipse Areas (SEA_C_) are depicted by a solid line with δ^13^C and δ^15^N values expressed in ‰. In A) both sea anemone hosts *Bartholomea annulata* and *Condylactis gigantea* were combined due to low samples sizes for *C. gigantea* (N = 2). Focal taxa are labeled by color.

**Fig. 4.**
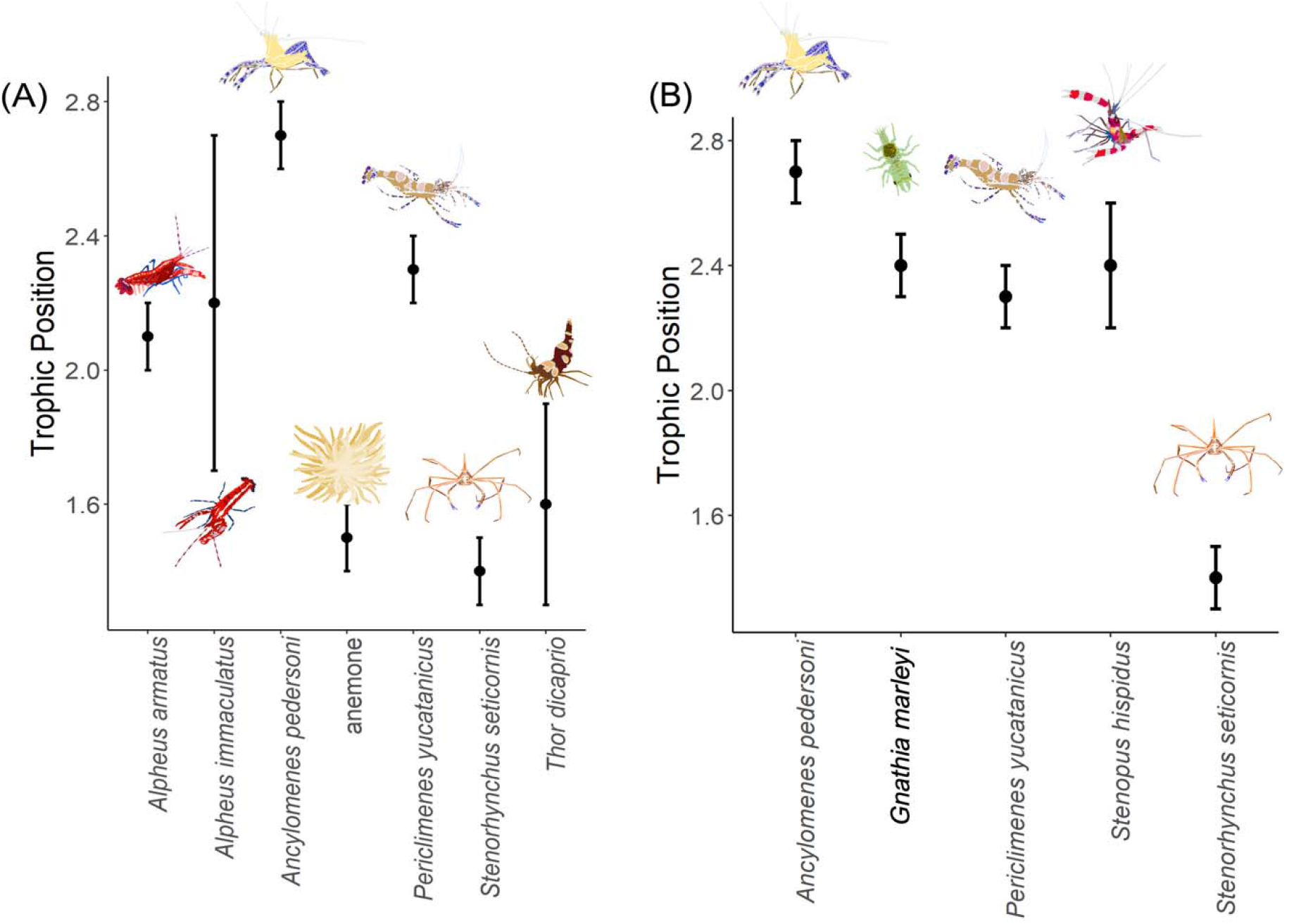
Mean and standard error calculations of trophic level for focal taxa within the A) sea anemone-crustacean symbiosis and B) putative crustacean cleaner species at St. Thomas, US Virgin Islands. In A) both sea anemone hosts *Bartholomea annulata* and *Condylactis gigantea* were combined due to low samples sizes for *C. gigantea* (N = 2).

Both cleaner shrimp species, *An. pedersoni* and *P. yucatanicus* occupied the highest trophic levels within the symbiosis but had significantly different isotopic niche spaces from one another (Fig. 3a, Table S1, Fig. S2). *Ancylomenes pedersoni* occupied the highest trophic level of any of our focal species and was significantly higher in δ^15^N than *P. yucatanicus* (p< 0.0001, see Table S2 for full statistical output for all linear models). Although both species exhibited the broadest isotopic niche spaces of our focal species, our analyses only calculated a 1% niche overlap between the species (Fig. 3a, Table S1). Among our co-occurring crustacean symbionts, we generally observe clearer partitioning of trophic level (i.e. δ^15^N) than δ^13^C values among co-occurring symbionts (Fig. 3a and 4a, Fig. S3). All pairwise comparisons of trophic level (δ^15^N) from our linear models were statistically significantly different at p < 0.0001 (Table S2). Only *S. seticornis* had δ^13^C ratios that were significantly different from all other taxa (Table S2, Fig. S3).

### Cleaner species niche partitioning

Among our dedicated and putative cleaner species, our data also reflect significant isotopic niche and trophic partitioning across all species (Fig. 3b and 4b, Table S2 and S3, Fig. S4 and S5). The dedicated cleaner *An. pedersoni* occupied the highest trophic level and a significantly different isotopic niche space from all other dedicated or putative cleaners (p < 0.0001, Fig. 3b and 4b, Table S2). Further, *An. pedersoni* occupied a significantly higher trophic level than the gnathiid isopod reef fish parasite (p < 0.0001, Fig. 3b and 4b). We calculated no more than a 5% isotopic niche space overlap between *An. pedersoni* and any other dedicated or putative cleaner (Fig. 3b, Table S3, Fig. S4). *P. yucatanicus*, currently considered a dedicated cleaner, occupied a significantly lower trophic level than *An. pedersoni*. Although its isotopic niche space and trophic level were significantly different than the putative cleaner *S. hispidus* (Fig. 3b and 4b, Table S2, Fig. S4), there was more isotopic niche overlap between these species (25%) than any other taxa (Table S3, Fig. S4). Both *P. yucatanicus* and *S. hispidus* occupied trophic levels similar to the gnathiid reef fish parasite (Fig. 3b and 4b). As above, *S. seticornis* remained an isotopic outlier from the other putative cleaners, occupying the lowest trophic level and exhibiting no niche overlap with any other species (Fig. 3b and 4b, Table S3, Fig. S4 and S5). Like our co-occurring anemone crustacean analyses, we observed more significantly partitioned trophic levels than δ^13^C values (Table S2, Fig. S5).

Comparisons within and between *An. pedersoni* and *P. yucatanicus* on a fine geographic scale show a significant effect of reef site on isotopic niche space and trophic level (Fig. 5, Table S2). Interestingly however, *An. pedersoni* and *P. yucatanicus* appear to be responding to reef site in different ways. At Brewers Bay *An. pedersoni* occupied a higher trophic level than at Flat Cay, whereas the trophic level of *P. yucatanicus* remained stable across reef sites (Fig. 5). In contrast, δ^13^C ratios do change in similar magnitude and direction across both species (Fig. 5) which may reflect the effect of local environmental conditions. These results suggest that environmental conditions alone cannot account for the change in δ^15^N observed in *An. pedersoni*.

**Fig. 5.**
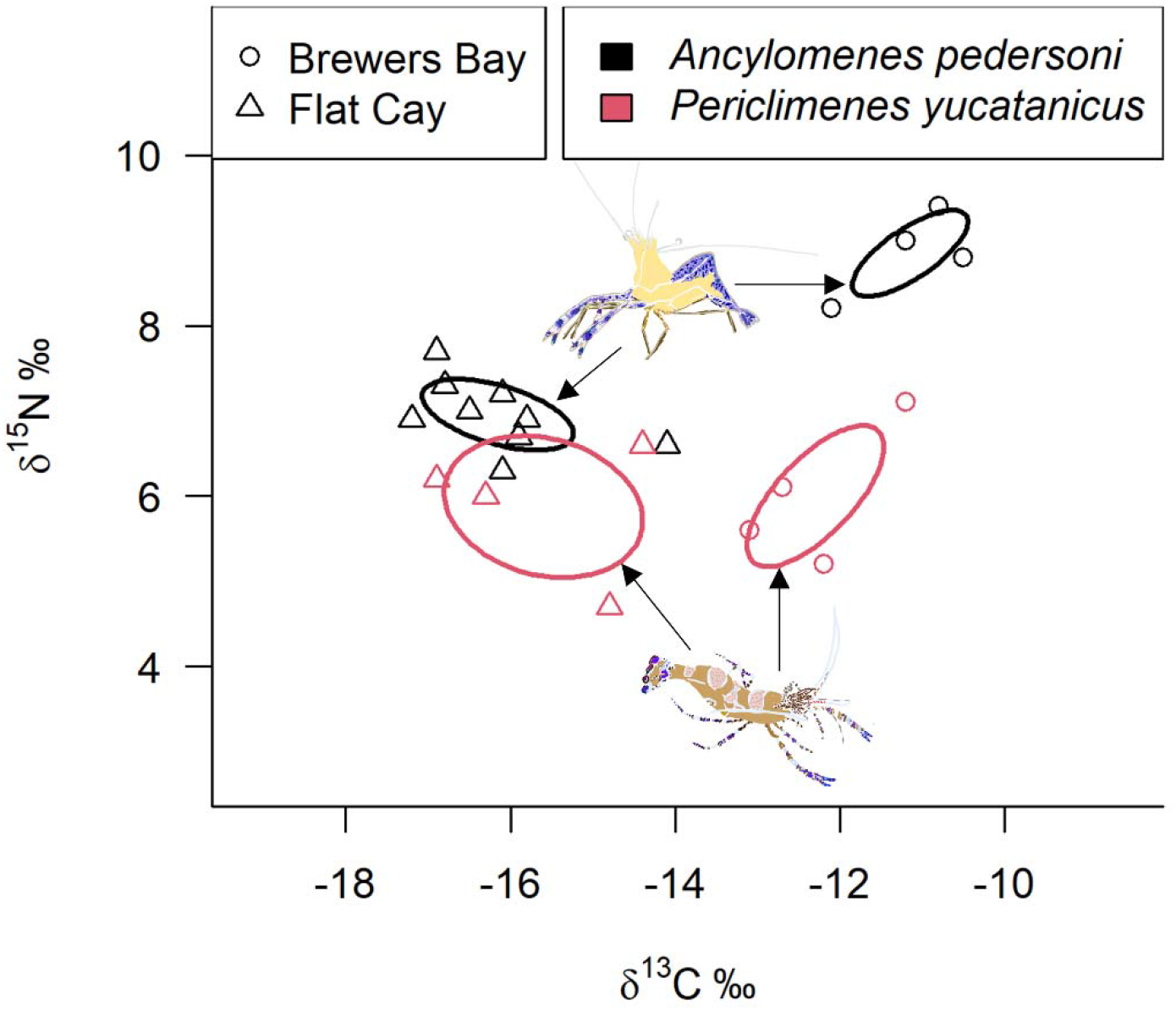
Isotopic Niche overlap for the cleaner shrimp *Ancylomenes pedersoni* (Black) and *Periclimenes yucatanicus* (Red) across two reef sites at St. Thomas, US Virgin Islands. 40% Standard Ellipse Areas (SEA_C_) are depicted by a solid line with δ^13^C and δ^15^N values expressed in ‰. Circles and triangles represent samples collected at Brewers Bay and Flat Cay, respectively.

## Discussion

The unified neutral theory of biodiversity and biogeography establishes a useful null hypothesis regarding niche partitioning and species co-occurrence because it presupposes that ecologically similar species that occur in sympatry are functionally analogous. Our data reject this null hypothesis in two ecological dimensions on a coral reef ecosystem in the Tropical Western Atlantic: 1) co-occurring crustacean symbionts of sea anemones that partition fine-scale microhabitat space around their hosts exhibit clear signatures of isotopic niche differentiation, and 2) dedicated and putative crustacean cleaner species exhibit isotopic niche differentiation that demonstrates these species differ in their functional roles as cleaners of reef fishes. Our data indicate that our focal taxa occupy distinct isotopic niche space and obtain food resources from significantly different sources, aligning well with the expectations of niche theory.

### Crustacean symbiont niche partitioning

Niche theory predicts that selection should drive co-occurring species into distinct ecological niche space to reduce interspecific competition and promote co-occurrence. Our *a priori* expectations (Table 1) were that crustacean symbionts that associated with *B. annulata* would fall into functional groups analogous to either detritivores or cleaners. As more species co-occur in a spatially restricted area than trophic levels and discernable ecological niches, we expected to recover isotopic signatures of functional redundancy within this symbiosis. Taken together, our data clearly reflect the opposite: co-occurring crustacean symbionts that partition fine-scale habitat space around *B. annulata* are partitioning ecological niche space as well and acquiring food resources in significantly different ways. Our stable isotope data aligns well with the expectations established by niche theory as each species occupies distinct isotopic niche space with minimal levels of overlap. Our data are especially well differentiated along the δ^15^N axis where we recover three clear partitions that correspond to different trophic levels. Both *T. dicaprio* and *S. seticornis* occupy low-level trophic levels that are aligned with our primary consumer mussel samples and the host anemone *B. annulata*- a species that is both photosymbiotic and heterotrophic. The snapping shrimps in the genus *Alpheus*, along with the spotted cleaner shrimp *P. yucatanicus*, occupy a higher trophic level analogous to secondary consumers (e.g. mantis shrimps). Pederson’s cleaner shrimp *An. pedersoni* occupies the highest trophic level, analogous to a high-level predator and consistent with expectations of a dedicated cleaner species that removes parasites from reef fishes.

The corkscrew sea anemone-crustacean symbiosis in the TWA has been a well-studied symbiotic system in relation to its population dynamics, population structure, phylogeography, symbiont diversity, distribution, and behavioral ecology (e.g. Briones-Fourzan et al. 2012, Huebner and Chadwick 2012a, b, O’Reilly et al. 2017, Titus et al. 2017a, 2019b, Caves et al. 2018, Titus and Daly 2017 2022). One recent major paper on the symbiosis demonstrated that the crustacean symbionts of *B. annulata* occur non-randomly across five unique microhabitat zones surrounding the host anemone. Interestingly, Huebner et al. (2019) found that although occasional overlap in microhabitat zones occurred between some species (mostly *An. pedersoni* and *P. yucatanicus*), each crustacean symbiont occupied a distinct microhabitat zone regardless of whether other crustacean symbionts were present. This consistent spatial microhabitat partitioning was hypothesized to limit competition for food among symbionts, but the diet and ecological niche of the crustacean symbionts were not disentangled. Our findings provide evidence in support of the hypothesis from Huebner et al. (2019) in that co-occurring species are not in competition for the same food resources.

Previous studies have variously described the trophic level and ecological function of our focal taxa qualitatively. As detailed here, some have been considered dedicated or putative cleaner organisms. Other studies have listed multiple members of this symbiosis as detritivores. Knowlton (1980) considered members of the *A. armatus* species complex to be detritivores based on behavioral observations, and Netchy et al. (2015) considered *S. seticornis* a detritivore in their survey of mobile invertebrate diversity along the Florida Reef Tract. Our data demonstrate that at our study sites in St. Thomas, members of the *A. armatus* species complex are more likely to rely on predation rather than detritivory given their broad trophic alignment with predatory mantis shrimps. We interpret the low trophic levels of *T. dicaprio* and *S. seticornis* to be a signature of heavy reliance on herbivory, although detritivory as a feeding strategy cannot be fully ruled out. While there was less differentiation along the δ^13^C axis among members of this symbiosis, *T. dicaprio* and *S. seticornis* were highly partitioned. We were unable to obtain planktonic samples to conduct mixing models to estimate the proportion of the diet obtained from pelagic vs. benthic carbon sources, but the depleted δ^13^C ratios of *T. dicaprio* suggest that this species may be more reliant on pelagic carbon sources while *S. seticornis* may be more reliant on benthic sources. Regardless, the non-overlapping isotopic niche space these taxa occupy indicate they are feeding on entirely different food resources. Given the tight linkage between the isotopic niche space of *T. dicaprio* and host anemone *B. annulata*, it is possible that both host and symbiont are relying on similar sources. It is important to note though that we had a low sample size for *T. dicaprio*, and in fact having a larger sample size may either increase or decrease the overlap between the two species.

Our data also shed light on the symbiotic status of many members within this crustacean community. In many host-symbiont interactions, it is often hypothesized or demonstrated that the host species is also a food resource for the symbionts (e.g. Fautin 1995, De Grave 2021), making the relationship more parasitic rather than commensal or mutualistic. Stable isotopes can partially disentangle these interactions without direct behavioral observation when the symbionts occupy trophic levels equal to, below, or multiple levels above the host. Highly differentiated δ^13^C signatures in a symbiont occupying a trophic level one step above the host would also be indicative of a mutualistic or commensal, rather than parasitic, interaction. Given these expectations, our dataset indicates that the host *B. annulata* does not serve as a significant food resource for *T. dicaprio*, *S. seticornis*, and *An. pedersoni*, as these species occupy either identical trophic levels to *B. annulata* with the same or different δ^13^C signatures or are multiple trophic levels above. These species can therefore be considered commensal or mutualistic with the host anemone *B. annulata*.

By contrast, our data cannot rule out that *B. annulata* serves as a potential food source for *P. yucatanicus* and members of the *A. armatus* species complex. Although we do not have direct isotopic fractionation calculations for these two species, fractionation was measured previously in *An. pedersoni* that were fed gnathiid isopod parasites (Demopoulous & Sikkel 2015), which was calculated at ∼1.4‰ and 1.3‰ for both δ^15^N and δ^13^C, respectively. If these values provide reasonable approximations for *P. yucatanicus* and the *A. armatus* complex, then *B. annulata* could be a potential food resource for both taxa. We are, however, dubious about the notion of a partially or fully parasitic relationship between the *A. armatus* complex and *B. annulata*. Both *A. armatus* and *A. immaculatus* are obligate symbionts to *B. annulata* and are well known to aggressively defend their host anemone from predatory fireworms (McCammon and Brooks 2014) as well as to rapidly unbury their host when covered in sediment (Perez-Botello et al. 2021). These mutualistic behaviors appear at odds with a feeding strategy that would impose significant fitness costs on their hosts, which we would expect to reduce the size of the host anemone and reduce the efficacy of the shelter members of the *A. armatus* species complex rely upon. No analogous behaviors have been observed for *P. yucatanicus*. Additional behavioral observations and possibly gut content metabarcoding would help disentangle this symbiotic relationship further. Stable isotope analyses from other invertebrate symbioses involving crustaceans and benthic macro-invertebrate hosts in Japan resolve a combination of commensal/mutualistic and parasitic relationships highlighting the diversity and complexity of these types of relationships (De Grave et al. 2021).

Given the ubiquity of symbiosis in nature, it is surprising how infrequently this framework is used to test alternative hypotheses surrounding species co-occurrence- especially on tropical coral reefs which are among the most conspicuously symbiotic ecosystems on the planet. From marine systems, most stable isotope analyses centered on symbiotic taxa are concerned with host-endosymbiont nutrient cycling (e.g. Ferrier-Pages and Leal 2018) or attempting to discern the symbiotic status of the host-symbiont relationship (i.e. whether it is mutualistic, commensal, or parasitic: De Grave et al. 2021, Puccinelli and McQuaid 2021). We are aware of only a small number of examples in the marine environment in which stable isotopes were used to disentangle isotopic niche partitioning and species co-occurrence within a host-symbiont framework. On coral reefs in southwest Madagascar, Terrana et al. (2019) demonstrated that palaemonid shrimp, gobiid fish, and polychaete worms that are co-occurring ectosymbionts of black corals and gorgonians exhibit a combination of isotopic niche overlap patterns. In this symbiosis, the coral host had significantly different isotopic niche space from all ectosymbionts, but there was significant overlap in trophic level between gobies and polychaete worms. However, the increased mobility of the symbiotic gobies over the polychaete worms likely allows for increased foraging opportunities, leading to the larger isotopic niche space observed for gobies and explaining species co-occurrence (Terrana et al. 2019). Both species of palaemonid shrimps exhibited significantly different isotopic niche space from each other and other guest symbionts. Terrana et al. (2019) concluded that in this system the host corals serve primarily as access to planktonic food resources in the water column, and that the coral hosts themselves are not significant sources of food for their ectosymbionts. Co-occurrence among symbionts is thus maintained via a combination of utilizing different food resources and niche breadth. In a mud-burrowing shrimp-bivalve commensalism where the burrowing shrimp provides habitat space for a co-occurring bivalve, species exhibited isotopic niche differentiation with the shrimp occupying a higher trophic level than the bivalve (Seike & Goto 2020). Finally, De Grave et al. (2021) also used stable isotopes to explore the trophic levels of symbiotic crustaceans and their benthic macro-invertebrate hosts from Japan, but like Seike & Goto (2020), this work used host-symbiont pairs rather than focusing on a community of species that co-occur on the same host. De Grave et al., (2020) found most host-symbiont pairs occupied different trophic levels. Generalizable conclusions about co-occurrence and niche partitioning within marine symbioses are difficult to synthesize given the lack of research in this area.

In terrestrial systems, analogous examples to our crustacean-sea anemone symbiosis are few, but the diverse symbioses insects form with host plants and other insect taxa are salient comparisons. In most cases, studies that employ stable isotopes to disentangle insect demonstrate, like ours, that co-occurring symbionts exhibit clear isotopic niche partitioning (e.g. Tillberg 2004, Parmentier et al 2016, Trimble and Sagers 2004, Sprenger et al. 2021). However, microhabitat partitioning by symbionts is not always indicative of trophic or isotopic niche differentiation. Stefan et al. (2015) demonstrated that co-occurring mite species that are symbiotic with seabirds exhibit strong microhabitat partitioning on bird feathers but are isotopically similar. The processes that allow for, and reinforce, species co-occurrence within symbioses are thus seldom discernable without integrative studies involving stable isotopes, behavioral observations, experimentation, and gut content analyses.

### Cleaner species niche partitioning

Cleaning interactions between client reef fishes and smaller cleaner organisms are among the most heavily studied behavioral interactions in the marine environment. Physical contact between client and cleaner is regularly used as a behavioral indication that a cleaning interaction has occurred (e.g. Huebner et al. 2012a, b, Titus et al. 2015a), which is presumed to result in the removal of a potentially harmful parasite from the client reef fish. Many cleaner organisms are also brightly colored and conspicuous on coral reef habitats, leading many closely related and similarly colored species to be labeled as cleaners as well (reviewed by Vaughan et al. 2017). The combination of behavior and color-pattern has created a low-bar for a species to obtain “cleaner” status. As a result, dozens of cleaners co-occur on the same reefs, and cleaning symbioses are expected to be among the most functionally redundant ecological interactions on coral reefs. Our data are the first to use stable isotopes to explore niche partitioning across a community of co-occurring species that have been labeled (using different standards of evidence) as cleaners and demonstrate high levels of niche partitioning and a distinct lack of functional redundancy among these species in the TWA. Only Pederson’s cleaner shrimp *An. pedersoni* occupies a trophic level that is consistent with expectations of a dedicated cleaner organism- a species that is a full-time cleaner that obtains most of its diet by removing parasites from reef fishes. Demopoulous and Sikkel (2015) calculated an δ^15^N isotopic fractionation for *An. pedersoni* that had been fed captive gnathiid isopods of ∼1.4‰. Our data are well aligned with this value and demonstrate that *An. pedersoni* sits a full trophic level above our gnathiid isopod parasite samples, the spotted cleaner shrimp *P. yucatanicus,* and the banded coral shrimp *S. hispidus*. Isotopically, *An. pedersoni* is thus a top-level predator on TWA coral reefs. From these data we conclude that *An. pedersoni* is the only dedicated cleaner shrimp species on coral reefs in the TWA.

Our results also provide important insight into the co-occurring spotted cleaner shrimp *P. yucatanicus* and begin to clarify long-standing ambiguity surrounding the ecological role of this species. Long considered a dedicated cleaner (Vaughan et al. 2017), our δ^15^N stable isotope data demonstrate that *P. yucatanicus* is clearly feeding at a lower trophic level than *An. pedersoni*. The δ^15^N values for *P. yucatanicus* are identical to our gnathiid isopod samples which indicates that reef fish parasites are not the overwhelming food resource for this species. As mentioned above, *P. yucatanicus* is a close relative of *An. pedersoni,* an obligate associate of sea anemones, and a species that have been observed engaging in behavioral interactions with reef fishes that are, by definition, “true” cleaning interactions (Titus et al. 2017a). Behavior alone, however, does not appear to be indicative of the ecological role of this species and we recommend this species no longer be described as a dedicated cleaner on TWA coral reefs.

What food resources *P. yucatanicus* primarily consumes and whether it removes parasites at all from reef fishes during their brief cleaning interactions is unclear. If *P. yucatanicus* is a facultative cleaner, e.g. removing the occasional parasite from reef fish but obtaining the majority of its food resources from lower trophic levels, the δ^15^N values we obtained would be consistent with this lifestyle. However, McCammon et al. (2015) showed *P. yucatanicus* did not significantly reduce monogenean flatworm parasites from reef fish, and our data here suggest gnathiids are not a major source of nutrition. Alternatively, *P. yucatanicus* could be ingesting fish mucus, scales, or other fish tissue rather than parasites during their brief interactions with posing client reef fishes. This hypothesis could imply that *P. yucatanicus* is a cleaner mimic species, where this species cheats posing clients by removing energetically expensive tissue rather than ectoparasites. Although an interesting hypothesis which has been proposed in the past (e.g. McCammon et al. 2010, Titus et al. 2017a), Demopoulous and Sikkel (2015) explored signatures of isotopic fractionation of gnathiid isopods that had been feeding on multiple species of reef fishes (e.g. French grunts, squirrelfish, and damselfish). Their findings demonstrate that gnathiids, which are only parasitic at the juvenile life stage, have δ^15^N values that are identical to their fish hosts. There is no isotopic fractionation of δ^15^N from host tissue to juvenile gnathiids until they digest their meal and metamorphose into non-parasitic adults. Thus, the juvenile gnathiid samples we included here that had been feeding on a variety of damselfish species should be interpreted as a direct proxy for the trophic level of their host. These were a mix of longfin, cocoa, and beaugregory damselfishes, which consume a diet of zooplankton, benthic algae, and invertebrates. This suggests that the food resources *P. yucatanicus* are ingesting are not coming from reef fishes that are secondary consumers and above, as any sort of resource acquisition derived from fish would place this species above gnathiid isopods and more in line with *An. pedersoni*. In this scenario, a cleaner mimic that cheats reef fishes would be isotopically indistinguishable from a dedicated cleaner species that feeds on gnathiids. A targeted study, specifically using gut content metabarcoding, is needed to provide more specific insight into *P. yucatanicus* diet than what stable isotopes can provide. What we can ascertain from our data is that *P. yucatanicus* is primarily ingesting animal tissue as it sits firmly within a secondary consumer trophic level, but we hypothesize these sources are not primarily derived from parasites or reef fishes.

Similar to *P. yucatanicus*, the δ^15^N stable isotope values we recovered for the banded coral shrimp *S. hispidus* also clearly reflects that this is not a dedicated cleaner shrimp species. Again, like *P. yucatanicus*, *S. hispidus* engages in behavioral interactions with reef fishes that involve physical contact which could indicate true cleaning interactions. This species however is not an anemone associate, but rather lives solitarily in crevices in coral reef substrate, and is occasionally observed cleaning moray eels and other nocturnal reef fishes that reside in dark sheltered habitats during the day (e.g. Limbaugh et al. 1961). The δ^15^N values for *S. hispidus* indicate that this species is primarily ingesting animal tissue, although the exact source remains unknown. Based on our data we suggest that at most, the banded coral shrimp be considered a facultative cleaner, but again, trophic level alignment with gnathiid isopods suggest that the majority of the food resources this species acquires are not derived from reef fishes. Both *P. yucatanicus* and *S. hispidus* do occupy the same general trophic level, and while we did calculate significant isotopic niche partitioning between these species, these had the most calculated isotopic overlap of any pairs of putative or dedicated crustacean cleaner species we studied here (25%). The primary difference between these taxa however, was that *P. yucatanicus* had a broader range of δ^13^C and thus, may have a broader ecological niche than *S. hispidus*. Additionally, both species occupy different microhabitats and thus there is minimally spatial partitioning between these taxa.

Unlike the above taxa, the δ^15^N values recovered for the yellowline arrow crab *S. seticornis* demonstrate that not only is this species not a dedicated or facultative cleaner species, but it likely does not ingest animal tissue at all at our field sites in St. Thomas, USVI. As a common associate of sea anemones in the TWA, any contact between *S. seticornis* and reef fish is likely incidental, and the result of a fish posing to be cleaned by *An. pedersoni*, which regularly co-occurs with *S. seticornis* at *B. annulata*. Although it is possible that *S. seticornis* takes on a different functional role in Brazil, where cleaning behavior for this species was first described, our data caution against this species being labeled a cleaner species in any capacity.

Finally, our fine-scale geographic sampling across reef sites provides interesting preliminary insight into the ecological function of both *An. pedersoni* and *P. yucatanicus,* as well as the role of the local environment on the stable isotope values of these species. For both, our data show a similar shift in magnitude in δ^13^C values for shrimp at Brewers Bay and Flat Cay, signifying that reef site may be largely responsible for driving the carbon signature at the base of the food chain, and that both species are responding in similar ways. At Flat Cay, a deeper offshore site with more oligotrophic conditions, depleted δ^13^C values may indicate a reliance on pelagic carbon sources. Shrimp from the nearshore Brewers Bay are more carbon enriched, which may indicate a reef site dominated by benthic primary sources. Interestingly however, δ^15^N values remain consistent for *P. yucatanicus* at both sites, which we interpret to mean that this species is acquiring food from the same resources at the same trophic level regardless of reef site. *An. pedersoni* on the other hand show significantly higher δ^15^N values at Brewers Bay than at Flat Cay. *An. pedersoni* is known to remove parasites from over 40 species and 20 families of reef fishes that encompass a wide range of herbivorous and predatory clients (e.g. Titus et al. 2015a, Romain et al. 2020). Thus, the diversity and trophic levels of the client community that visits *An. pedersoni* could have a significant impact on shrimp δ^15^N values if fish communities differ between reef sites. Although this finding should be considered preliminary and deserves more detailed investigation, this pattern is consistent with the interpretation that *P. yucatanicus* is not a dedicated cleaner shrimp nor functionally equivalent with *A. pedersoni*. Ecological differences between Brewer’s Bay and Flat Cay appear to result in changes in the observed isotopic trophic level of *An. pedersoni* without a concomitant change in *P. yucatanicus*.

Our stable isotope data represent the first comparative study of co-occurring cleaner species from a coral reef ecosystem. We show that *An. pedersoni* is not only the most ecologically important crustacean cleaner species on TWA coral reefs, but likely the *only* significant crustacean cleaner species on TWA coral reefs. Future research should extend our comparative study into the cleaner fish community by comparing *A. pedersoni* with the co- occurring neon cleaner goby *Elacatinus oceanops*. Titus et al. (2015a) calculated extensive overlap in the reef fish client communities that visit both cleaner species but highlighted important differences in the duration of the cleaning interactions, with *An. pedersoni* providing significantly longer cleans than *E. oceanops*. Whether these cleaners are targeting the same parasite communities on client reef fish, and thus, potentially competing for resources and providing the same functionally redundant services to reef fishes, remains a major unresolved question in the cleaning literature and would shed light on the ultimate and proximate causes of cleaning seeking behavior in reef fishes (Titus et al. 2015a).

### Speculations

The cleaning symbiosis literature is rich in examples demonstrating that cleaner species make conscious decisions regarding the services they provide to client reef fishes. These can include providing higher quality services to predatory clients, preferring large over small clients, and simply choosing not to engage in cleaning interactions with posing client fishes. For dedicated cleaners, these decisions will impact the source and isotopic makeup of the parasites they ingest, and by extension, the isotopic material they accumulate. Our fine-scale geographic data show that *An. pedersoni* are foraging at higher trophic levels at Brewers Bay than at Flat Cay, but only four of our samples came from Brewers Bay. Are *An. pedersoni* at Brewers Bay choosing to engage more with predatory clients than at Flay Cay, or is the client community composition at Brewers Bay skewed towards predatory reef fishes? Does cleaner shrimp social group hierarchy impact cleaner access to specific clients? Does social status, behavior, and client diversity interact to drive variation in individual cleaner shrimp isotopic content? If so, our data may imply a useful role for stable isotopes in disentangling individual variation between cleaners and the clients they clean, and ultimately, capture social and behavioral status of cleaner organisms with a single data point.

### Conclusions

When ecologically similar species co-occur, it creates difficult conceptual problems to disentangle in ecology. On one hand, functional redundancy is considered a hallmark of resilient ecological communities, particularly in highly biodiverse ecosystems such as coral reefs (e.g. reviewed by Biggs et al. 2020). Communities with co-occurring, functionally redundant species are expected to be better able to buffer disturbance events while maintaining ecosystem function. On the other hand, functional redundancy appears to be at odds with stable species co-existence in the long-term (e.g. Loreau 2004). Our data demonstrate little, if any, evidence of functional redundancy both within the community of crustacean symbionts that co-occur mutualistically with the corkscrew sea anemone *B. annulata*, or among a community of co-occurring species variously described as cleaner species in the TWA. These species are clearly partitioning isotopic niche space, acquiring resources in different ways, and providing different ecological services to the broader community. The lack of functional redundancy in the cleaner shrimp community is particularly interesting but potentially concerning. Like corals, sea anemones bleach in response to prolonged exposure to high water temperatures. As *An. pedersoni* is an obligate symbiont of sea anemones and the only dedicated cleaner shrimp in the TWA, bleaching events will eliminate cleaning stations and may have dramatic radiating impacts across the entire community of reef fishes if there is a significant reduction in access to cleaners.

Finally, although stable isotope studies on co-occurring free-living organisms abound, and have been performed from every major ecosystem on earth, far less research has been focused on guilds of symbiotic taxa that co-occur on a single host and form an assemblage of species. Mutualisms are among the most important ecological interactions on the planet and have contributed enormously to global levels of biodiversity. Studies focused explicitly on mutualism and species co-occurrence will thus contribute a more holistic understanding of not only how symbiosis has generated biodiversity, but how it is maintained.

