## Supplemental Material for "Stable isotopes disentangle niche partitioning and species co-occurrence in a multi-level marine mutualism"

**Supplementary Tables**

**Table S1**. Results of pairwise percent niche overlap (gray, top right) and isotopic area overlap (blue, bottom left) for co-occurring species within the sea anemone-crustacean symbiosis at St. Thomas, US Virgin Islands. The spotted cleaner shrimp *Periclimenes yucatanicus* is abbreviated *P. yuc* for space, as is the pistol snapping shrimp *Alpheus immaculatus* (*A. imm*)*.*

|  |  | **Percent Niche Overlap (%)** | | | | | | |
| --- | --- | --- | --- | --- | --- | --- | --- | --- |
|  |  | *A. pedersoni* | *P. yuc* | *T. dicaprio* | *A. imm* | Sea anemone | *A. armatus* | *S. seticornis* |
| **Overlap Niche Area (%_o_^2^)** | *A. pedersoni* |  | 1% | 0% | 0% | 0% | 0% | 0% |
|  | *P. yuc* | 0.11 |  | 0% | 9% | 0% | 14% | 0% |
|  | *T. dicaprio* | 0 | 0 |  | 0% | 18% | 0% | 0% |
|  | *A. imm* | 0 | 0.46 | 0 |  | 0% | 20% | 0% |
|  | Sea anemone | 0 | 0 | 0.47 | 0 |  | 0% | 0% |
|  | *A. armatus* | 0 | 0.77 | 0 | 0.33 | 0 |  | 0% |
|  | *S. seticornis* | 0 | 0 | 0 | 0 | 0 | 0 |  |

**Table S2.** Full statistical output for all linear models conducted for all analyses. **Bold** text under Response Variable highlights the specific dataset analyzed.

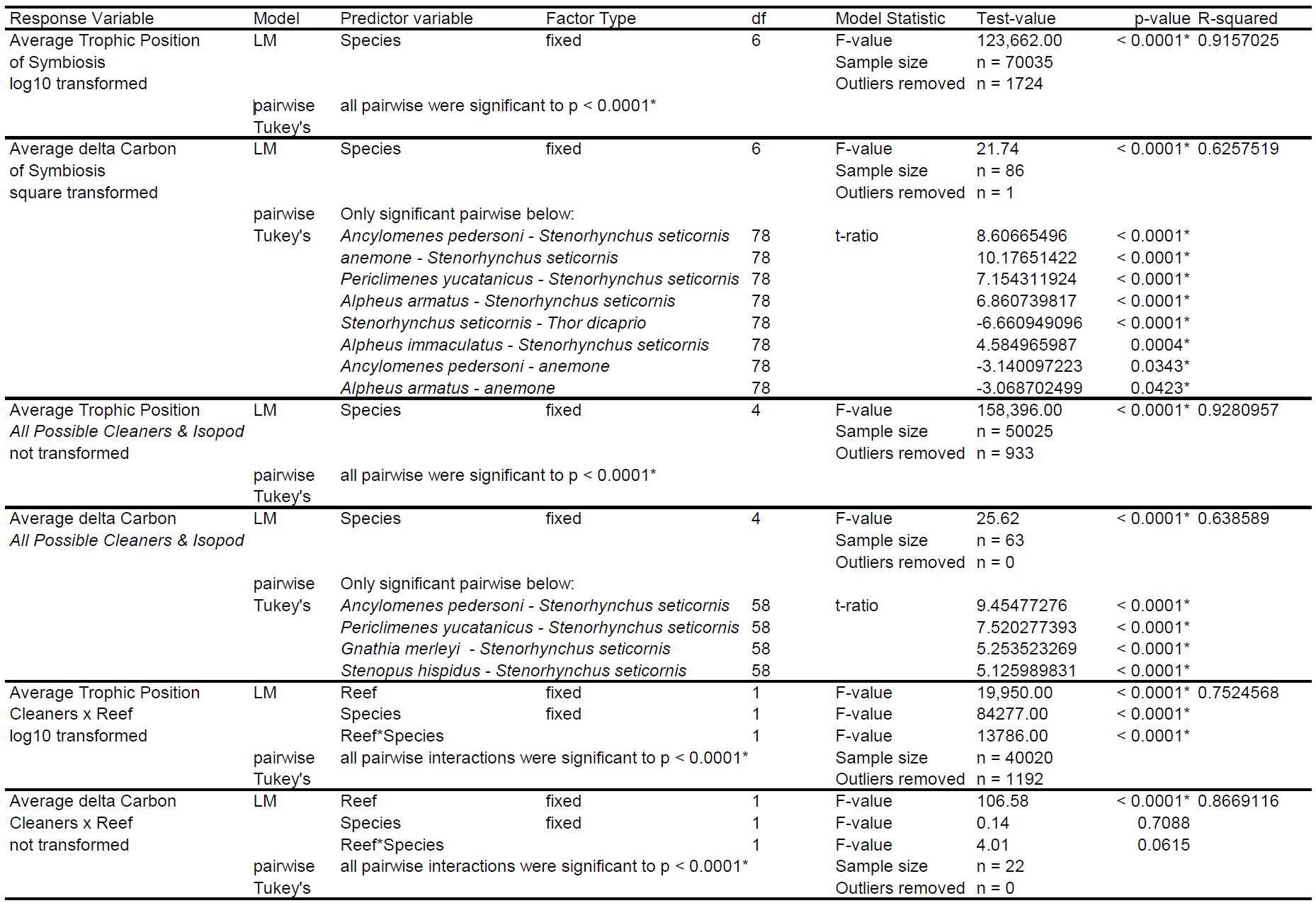

**Table S3**. Results of pairwise percent niche overlap (gray, top right) and isotopic area overlap (blue, bottom left) for dedicated and putative cleaner species at St. Thomas, US Virgin Islands. *Gnathia marleyi* represents a common reef fish gnathiid isopod parasite included in the analysis as a potential food source for all focal taxa.

|  |  | **Percent Niche Overlap (%)** | | | | |
| --- | --- | --- | --- | --- | --- | --- |
| **Overlap Niche Area (%_o_^2^)** |  | *A. pedersoni* | *P. yucatanicus* | *G. marleyi* | *S. seticornis* | *S. hispidus* |
|  | *A. pedersoni* |  | 1% | 0% | 0% | 5% |
|  | *P. yucatanicus* | 0.07 |  | 7% | 9% | 25% |
|  | *Gnathia marleyi* | 0 | 0.36 |  | 0% | 8% |
|  | *S. seticornis* | 0 | 0 | 0 |  | 0% |
|  | *S. hispidus* | 0.33 | 1.48 | 0.20 | 0 |  |

**Supplementary Figures**

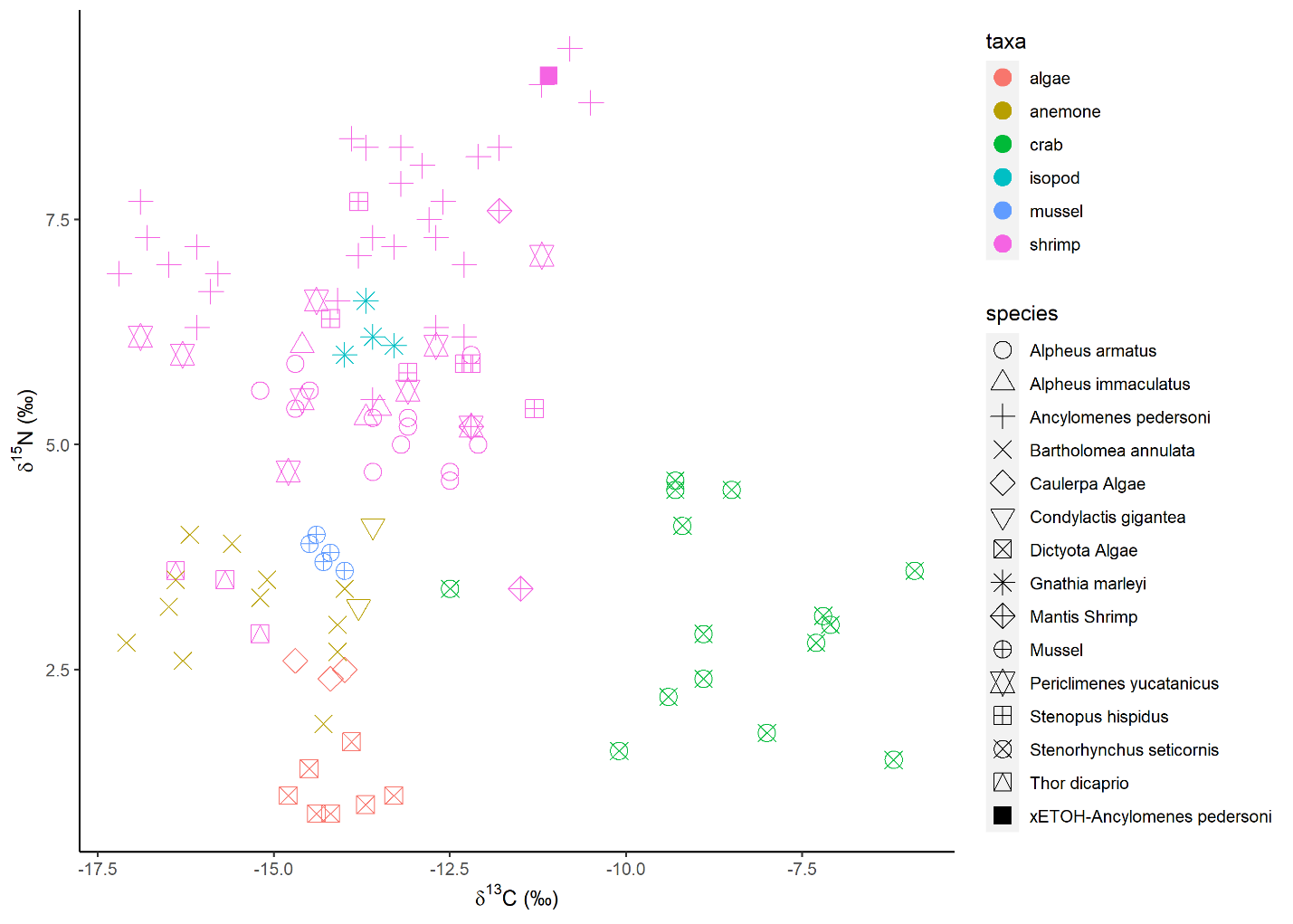

**Figure S1**. Isotopic biplot of for δ^13^C and δ^15^N ratios for all samples plotted individually. Species are labeled by shape and color coded by broad taxonomic group.

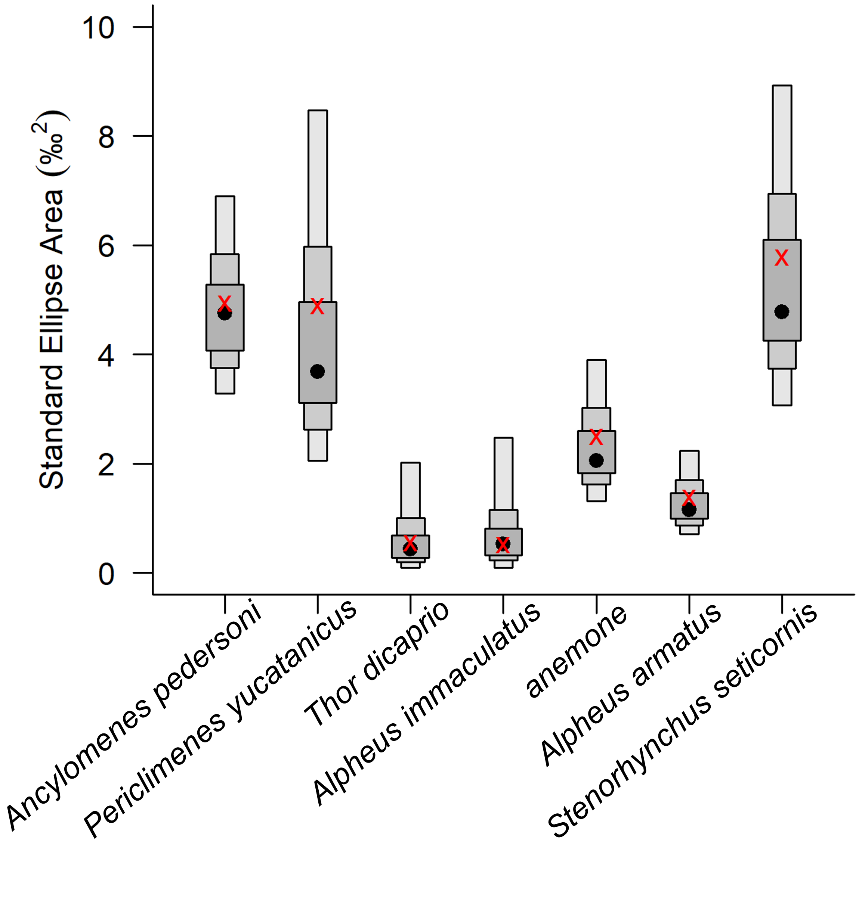

**Figure S2**. Standard ellipse area (SEA_C_) overlap analysis for the sea anemone-crustacean focal taxa. Data are corrected for low sample size (red crosses) and reflect the mode (black dot) and 99%, 95%, 50% intervals.

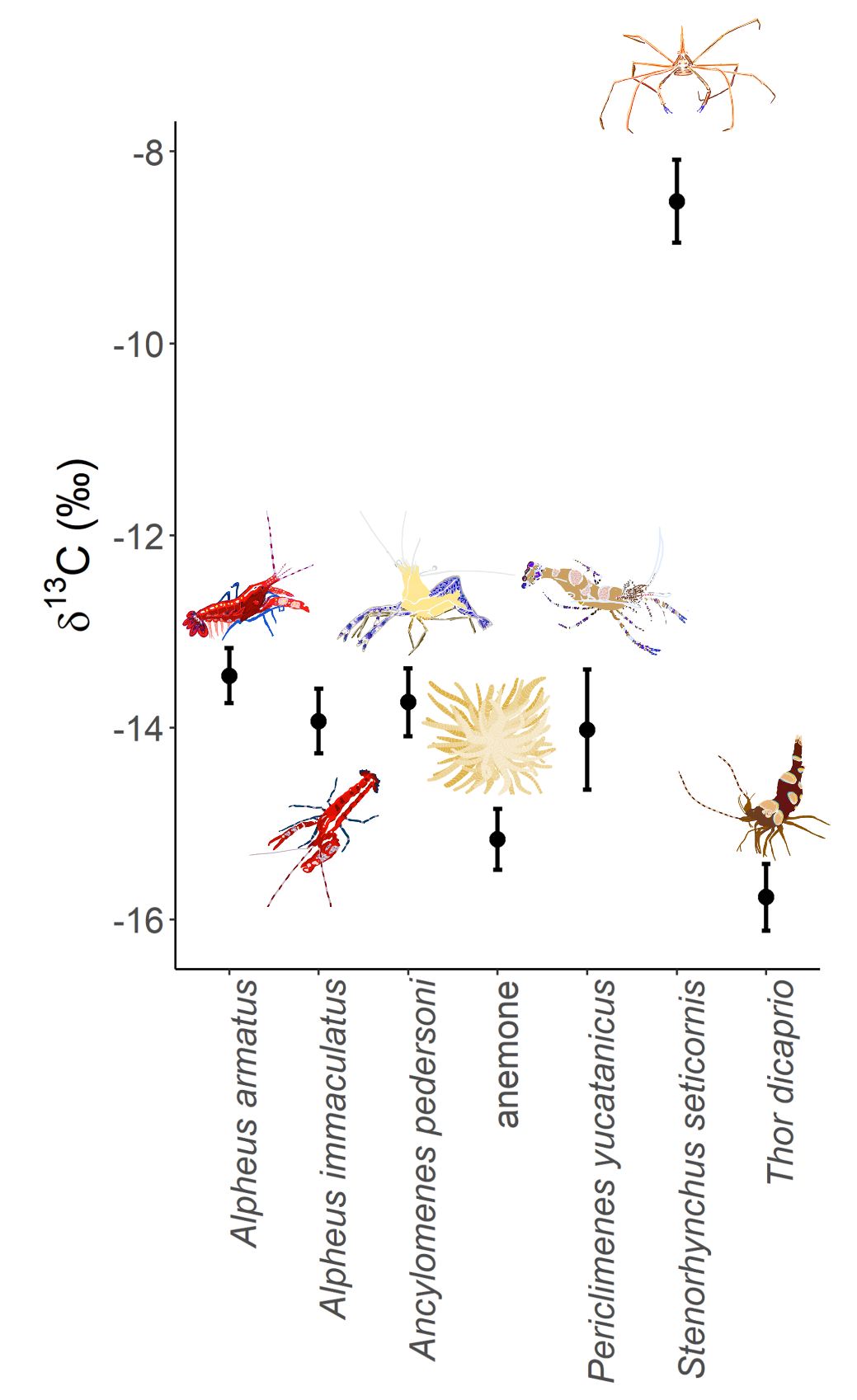

**Figure S3.** Mean and standard error of δ^13^C for all focal taxa within the sea anemone crustacean symbiosis at St. Thomas, US Virgin Islands.

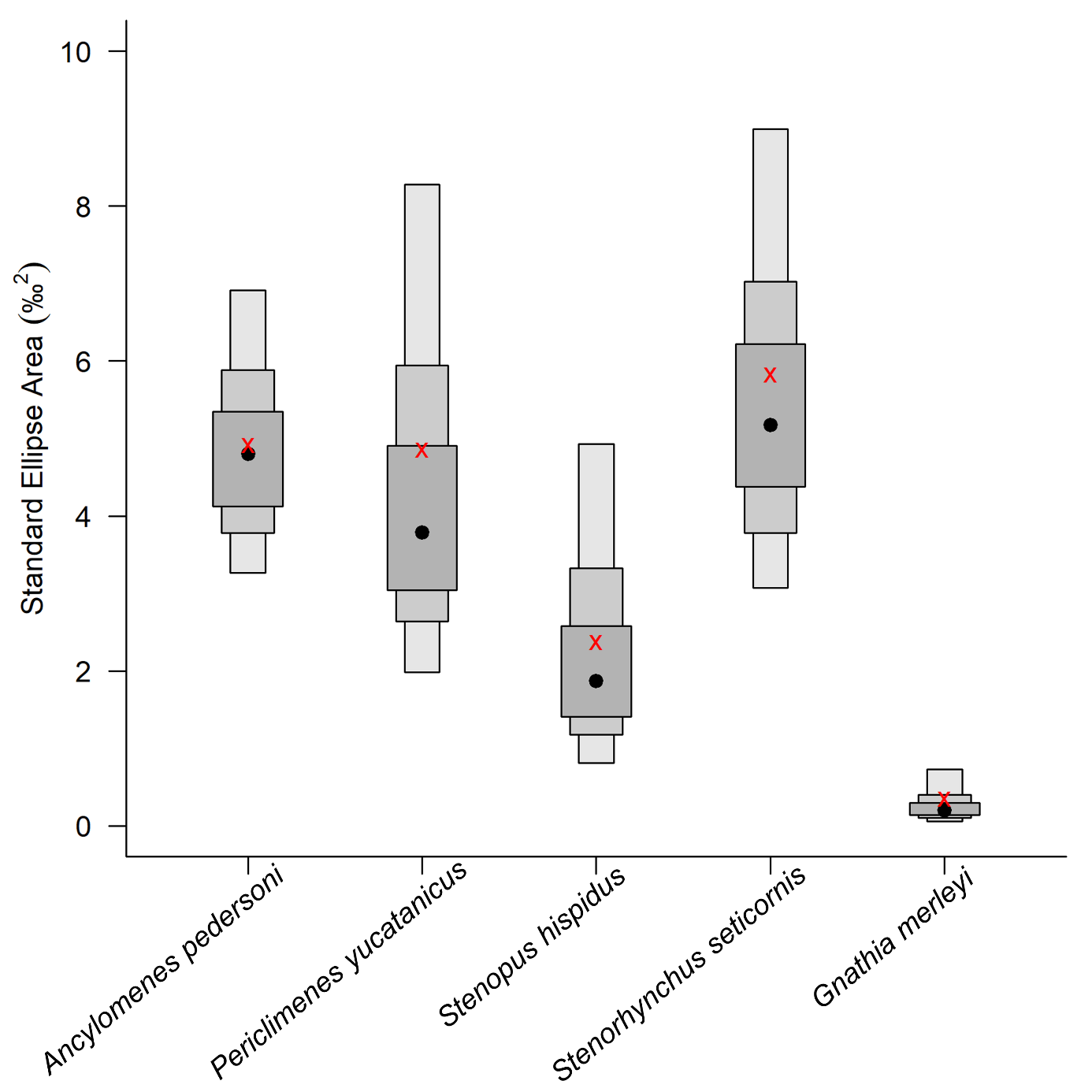

**Figure S4.** Standard ellipse area (SEA_C_) overlap analysis for the dedicated and putative cleaner species. Data are corrected for low sample size (red crosses) and reflect the mode (black dot) and 99%, 95%, 50% intervals.

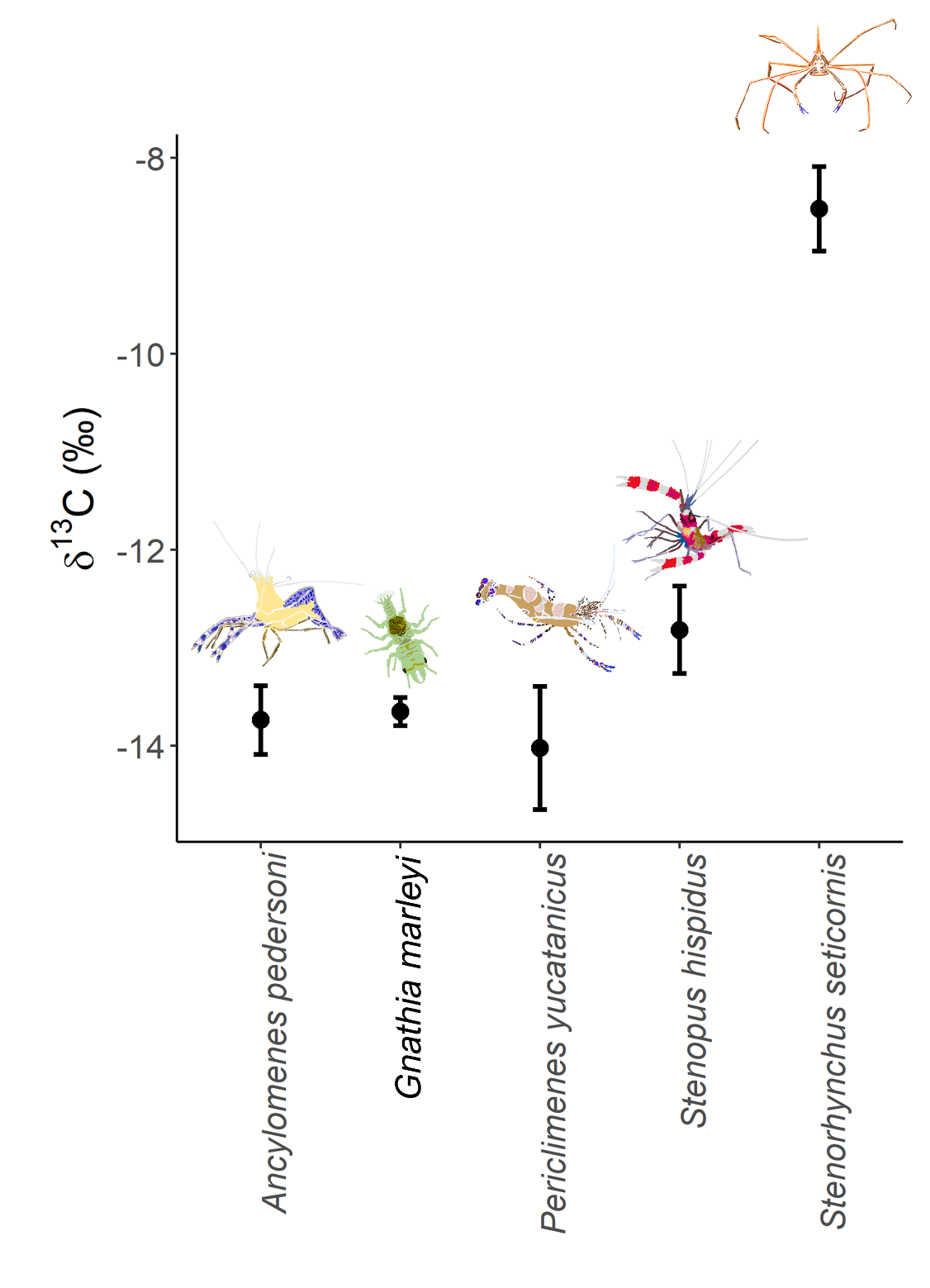

**Figure S5.** Mean and standard error of δ^13^C for dedicated and putative cleaner species at St. Thomas, US Virgin Islands

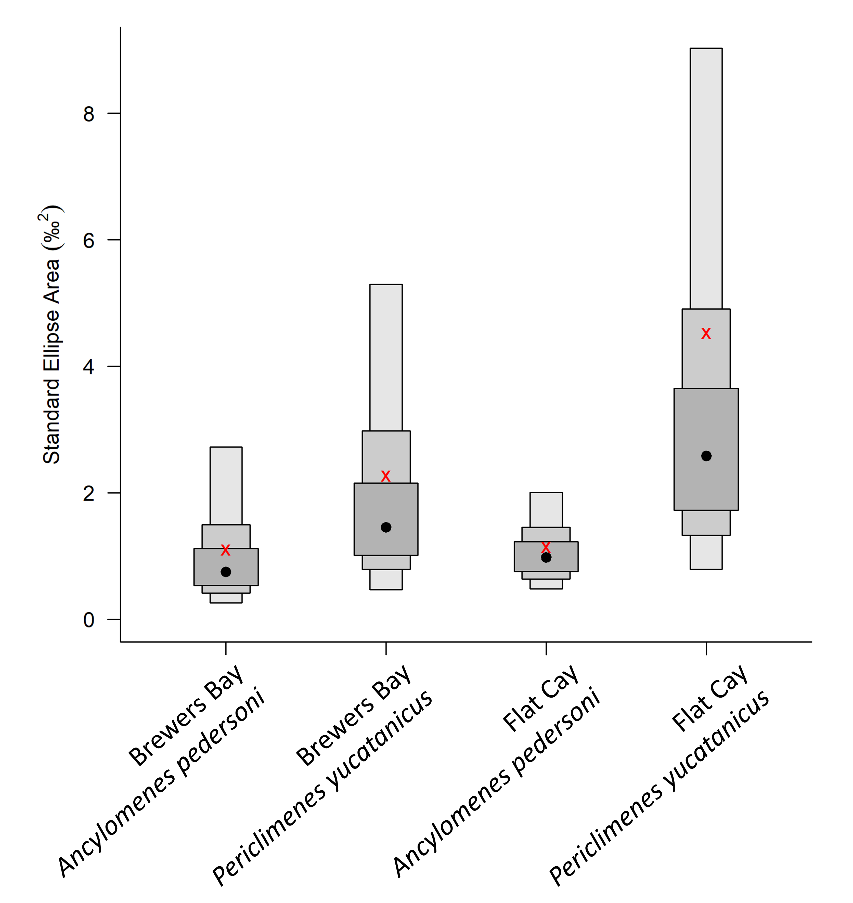

**Figure S6**. Standard ellipse area (SEA_C_) overlap analysis for the dedicated cleaner species *Ancylomenes pedersoni* and *P. yucatanicus* between reef sites at St. Thomas, US Virgin Islands. Data are corrected for low sample size (red crosses) and reflect the mode (black dot) and 99%, 95%, 50% intervals.
